# Molecular Architecture of *Cryptococcus* Cell Walls Reveals Species-Specific Chitosan-Dependent Remodeling

**DOI:** 10.64898/2026.04.14.718415

**Authors:** Ankur Ankur, Rajendra Upadhya, Mahsa Doosti, Davis Ferreira, Li Xie, Ivan Hung, Jennifer K. Lodge, Tuo Wang

## Abstract

*Cryptococcus neoformans* and *Cryptococcus gattii* are fungal pathogens that cause life-threatening infections, including cryptococcal meningitis. A distinctive feature of the cryptococcal cell wall is the extensive deacetylation of chitin to chitosan, a modification that is essential for virulence but whose structural role in cell-wall organization remains poorly understood. Here, we analyzed the cell walls of wild-type strains of both species and their avirulent chitosan-deficient mutants, which serve as vaccine candidates. Loss of chitosan disrupted cell morphology and altered cell-wall ultrastructure, with more pronounced defects in *C. neoformans*. Solid-state NMR revealed that aggregated α-1,3-glucans form the principal rigid domain of the cell wall in both species and are closely associated with chitin microfibrils, whereas surrounding β-glucans and mannoproteins constitute a more dynamic matrix. Chitosan modulates hydration and flexibility, and its loss increases chitin exposure and triggers species-specific remodeling of the polysaccharide network. In *C. neoformans*, chitosan depletion increased α-1,3-glucan content and reduced β-glucan levels, whereas *C. gattii* selectively lost one α-1,3-glucan subtype while maintaining β-glucan levels. Although capsule production remained intact, chitosan deficiency altered glucuronoxylomannan linkage patterns and mannoprotein composition. These findings reveal how chitosan organizes cryptococcal cell-wall architecture and highlight distinct structural adaptation strategies among pathogenic *Cryptococcus* species.

## INTRODUCTION

The *C. neoformans* and *C. gattii* species complexes are the causative agents of cryptococcosis, a life-threatening fungal disease that primarily manifests as meningoencephalitis in humans^1,2^. The *C. neoformans* species complex is considered an opportunistic pathogen because it predominantly infects immunocompromised individuals, particularly those with HIV/AIDS^3,4^. In contrast, the *C. gattii* species complex is regarded as a primary pathogen due to its ability to cause disease in otherwise immunocompetent hosts^4-6^. Cryptococcal meningitis affects approximately 200,000 individuals worldwide each year and is associated with mortality rates ranging from 20-70%, depending on disease severity and host condition^7,8^.

Current treatment for cryptococcosis relies on prolonged combination therapy with amphotericin B, flucytosine, and fluconazole, a regimen that is both costly and associated with substantial toxicity^9-12^. Although echinocandins, antifungal drugs that inhibit β-1,3-glucan synthesis, are effective against many fungal pathogens, they exhibit limited activity against *Cryptococcus*, largely due to the distinctive architecture of the cryptococcal cell wall^13-16^. Consequently, clinical outcomes remain constrained by drug toxicity, emerging resistance, and persistently high mortality, emphasizing the need for improved therapeutic and preventative strategies, including effective vaccines.

A unique feature of the cryptococcal cell wall is the extensive conversion of chitin to chitosan through the action of chitin deacetylases^17,18^. This modification is essential for maintaining cell-wall integrity and virulence^19^. Chitosan levels can be altered either genetically, by deleting genes required for its biosynthesis, or environmentally, for example by culturing cells in unbuffered minimal medium^20^. In both cases, chitosan depletion results in strikingly similar phenotypes, including weakened cell-wall integrity, altered wall architecture, and reduced virulence in murine infection models^19,21,22^. Fluorescence labeling with cell-wall probes such as calcofluor white, wheat germ agglutinin, concanavalin A, and β-1,3-glucan-specific antibodies has demonstrated increased accessibility of multiple wall components in chitosan-deficient cells, suggesting that loss of chitosan reorganizes the wall and exposes pathogen-associated molecular patterns (PAMPs)^20^.

The importance of chitosan is further highlighted by the avirulent phenotype of the *cda1*Δ*2*Δ*3*Δ mutant, generated by deletion of the three chitin deacetylase genes^23^. In a murine inhalation model, this mutant is rapidly cleared from the lungs and fails to establish infection, owing to an altered host immune response compared with that induced by the wild-type strain^24^. Pulmonary vaccination with live *cda1*Δ*2*Δ*3*Δ cells confers robust protection against subsequent lethal challenge with virulent *C. neoformans*, and even heat-killed cells induce significant protective immunity^24^. In contrast, although deletion of the same three CDA genes in the hypervirulent *C. gattii* R265 strain also produces an avirulent mutant, vaccination with these cells fails to generate comparable protection against subsequent infection with wild-type R265^23^. These divergent outcomes suggest fundamental differences in how the two species interact with host immunity, potentially arising from species-specific differences in cell-wall composition and architecture.

Consistent with this idea, numerous studies have reported significant differences in pathogenic mechanisms between *C. neoformans* and *C. gattii*. Epidemiologically, *C. neoformans* infections occur predominantly in immunocompromised hosts, whereas *C. gattii* frequently infects immunocompetent individuals^23^. At the molecular level, *C. gattii* synthesizes substantially more chitosan than *C. neoformans* under both laboratory and in vivo conditions, and the regulation of chitosan biosynthesis differs between the species: in *C. neoformans*, Cda1 and Cda2 coordinately control chitosan production during infection, whereas in *C. gattii* R265, Cda3 plays the dominant role^23,25,26^. These observations suggest that species-specific differences in cell-wall organization may influence immune recognition and disease outcome. Notably, R265 has been shown to evade the host immune response more efficiently than KN99 in mice vaccinated with *cda1*Δ*2*Δ*3*Δ cells and subsequently coinfected with a KN99 and R265^27^.

Despite the central role of the fungal cell wall in virulence and immune interactions, its supramolecular architecture remains incompletely understood. Solid-state NMR spectroscopy has recently emerged as a powerful tool for high-resolution characterization of polysaccharides in intact fungal cells, enabling direct analysis of molecular organization and interactions within the native cell wall without disruptive extraction procedures^28-34^. A recent solid-state NMR study of *C. neoformans* has identified multiple isoforms of α-1,3-glucan and revealed a hierarchical organization of cell-wall polymers, in which rigid α-glucan domains associate with chitin, chitosan, capsule components, and melanin, while the mobile matrix is dominated by β-1,6-glucan with minor contributions from β-1,3-glucan^35^.

Here we extend this approach to compare the cell-wall architecture of *C. neoformans* KN99 and *C. gattii* R265 and their respective *cda1*Δ*2*Δ*3*Δ mutants. By combining targeted gene deletion with multidimensional ^13^C solid-state NMR spectroscopy, fluorescence-based cell-wall probes, and electron microscopy, we demonstrate substantial structural differences between the two species. Our analysis shows that although α-1,3-glucans form the dominant rigid domains in both organisms, chitin, chitosan, and β-1,6-glucan contribute more prominently to maintaining overall cell-wall integrity in *C. gattii*. These two species exhibit fundamentally distinct remodeling responses to chitosan depletion. Such species-specific strategies likely influence immune recognition and may help explain differences in pathogenicity between cryptococcal pathogens.

## RESULTS

### Chitosan is essential for maintaining proper cryptococcal cell wall architecture

To establish the cellular context for subsequent solid-state NMR analysis, fungal cultures were first examined by light and electron microscopy to assess cell morphology and cell wall ultrastructure. Light microscopy showed that wild-type strains of both species exhibited the typical round yeast morphology with normal budding and well-separated cells of relatively uniform size (**Fig. 1a**). In contrast, the chitosan-deficient *cda1*Δ*2*Δ3Δ mutants displayed aberrant morphology characterized by heterogeneous cell sizes and frequent clumping with reduced budding. These defects were more pronounced in the mutant derived from the *C. neoformans* KN99 background than in the corresponding *C. gattii* R265 mutant.

**Figure 1.**
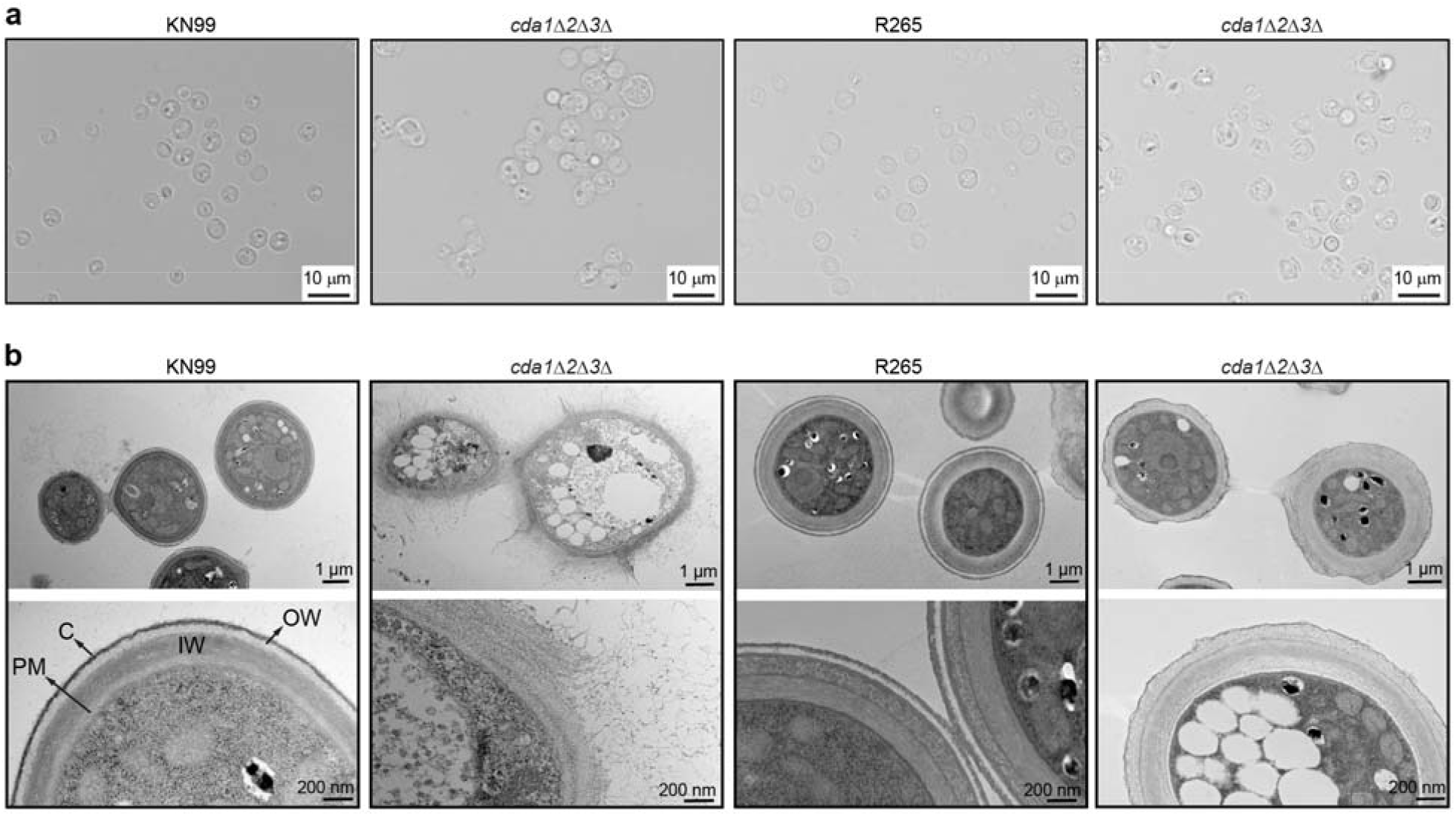
Cryptococcal cell wall architecture is severely altered in the absence of chitosan. (**a**) Bright field images of the wild-type and *cda1*Δ*2*Δ*3*Δ mutants of KN99 and *C. gattii* R265 background that were subjected to solid-state NMR analysis. (**b**) Representative transmission electron micrographs of cells obtained from the same cultures used for NMR analysis. From left to right, the four columns represent KN99 and its *cda1*Δ*2*Δ*3*Δ mutant, followed by R265 and its *cda1*Δ*2*Δ*3*Δ mutant. Following fixation, samples were processed, embedded, ultrathin_sectioned, and imaged by transmission electron microscopy. The inner wall (IW), outer wall (OW), plasma membrane (PM), and capsule (C) can be distinguished.

Transmission electron microscopy (TEM) further revealed pronounced differences in cell wall organization between the two species **(Fig. 1b)**. Wild-type cells of both species exhibited well-defined inner and outer cell wall layers, an intact plasma membrane, and an external capsular layer. Notably, wild-type *C. gattii* R265 cells displayed increased stratification of the cell wall. This enhanced stratification may result from increased staining of the chitosan-rich wall, as protonated amino groups in chitosan favor retention of the anions present in uranyl acetate, or from altered interactions of the stain due to the elevated chitosan content. Two distinct layers within the inner cell wall were clearly resolved in *C. gattii* but were not observed in *C. neoformans*, indicating a fundamental structural difference between the cell walls of the two species.

Consistent with the species-specific differences observed in wild-type cells, chitosan-deficient *cda1*Δ*2*Δ*3*Δ mutants also displayed marked differences in cell wall architecture. The *cda1*Δ*2*Δ*3*Δ mutant of *C. gattii* largely retained an organized cell wall structure with discrete inner and outer layers. In contrast, the *cda1*Δ*2*Δ*3*Δ mutant of *C. neoformans* exhibited a diffuse, poorly organized cell wall characterized by a “goopy” appearance, frequent detachment of the outer wall layer, and a wrinkled plasma membrane with prominent cytoplasmic invaginations (marked by asterisks).

### Chitosan depletion is accompanied by the removal of a α-1,3-glucan subtype in *C. gattii*

In both *C. neoformans* and *C. gattii*, chitosan was primarily observed in the rigid domains of the cell wall rather than the soft matrix, suggesting an important structural role of this molecule in maintaining wall strength. Rigid polysaccharides were analyzed using a cross-polarization (CP)-based 2D ^13^C-^13^C CORD correlation experiment^36^. Two distinct chitosan subtypes, designated Cs^a^ and Cs^b^, were unambiguously distinguished by their C1-C2 cross-peaks (Cs1-2) at (102.8, 56.7) ppm and (98.8, 56.8) ppm, respectively (**Fig. 2a, b**). These subtypes may arise from local structural differences in hydrogen-bonding patterns and molecular conformation and were completely absent in the corresponding chitosan-deficient mutants (**Fig. 2a**).

**Figure 2.**
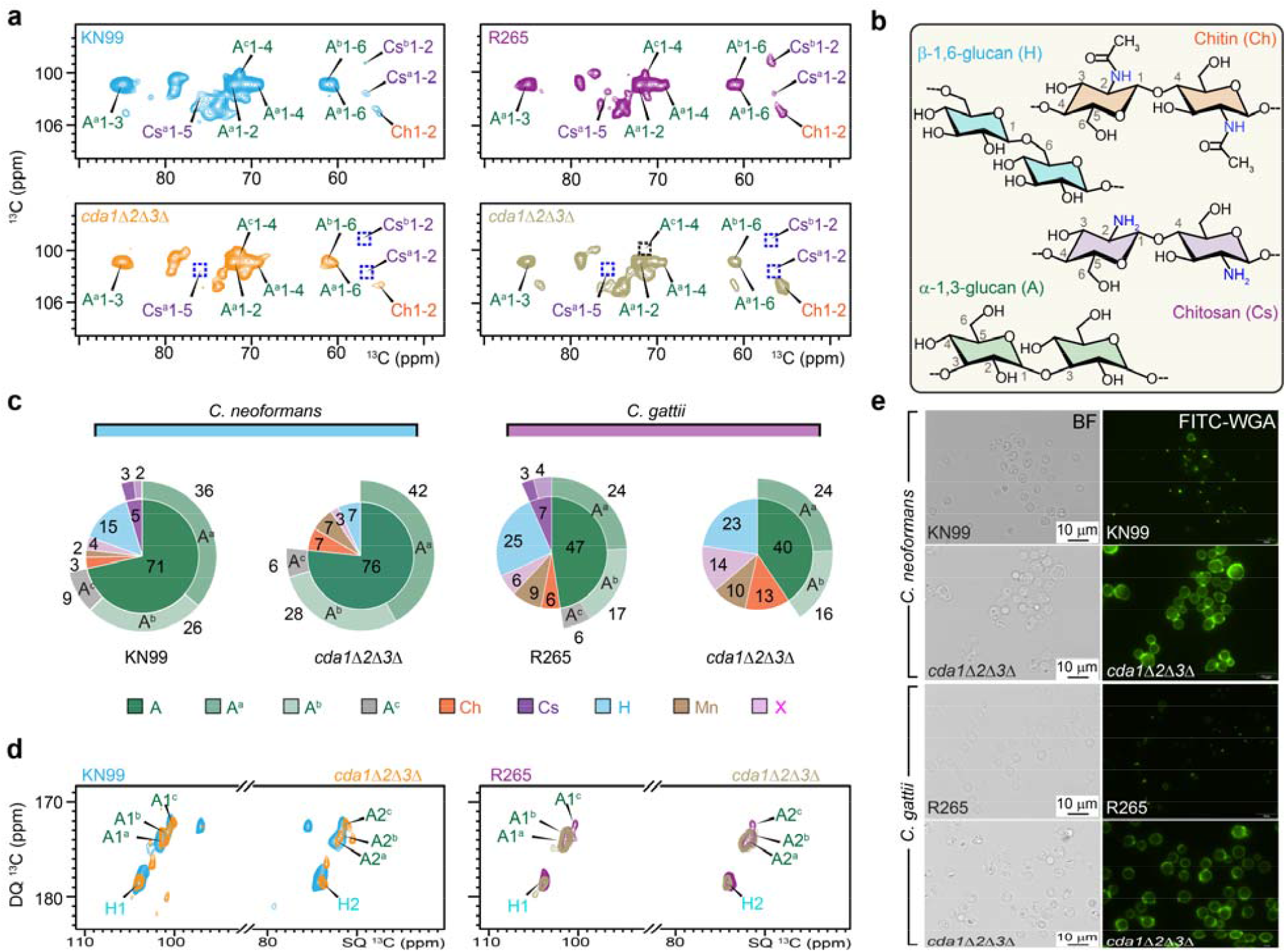
Altered rigid polysaccharides in wildtype and mutant strains of *C. neoformans* and *C. gattii*. (**a**) 2D ^13^C-^13^C CORD correlation spectra of *C. neoformans* and *C. gattii* cells. The absence of chitosan in *cda1*Δ*2*Δ*3*Δ mutants of each species is highlighted with dashed blue squares. The absence of type-c α-1,3-glucan signal in *C. gattii cda1*Δ*2*Δ*3*Δ mutant is highlighted with dashed black square. Observed cell wall carbohydrates include α-1,3-glucan (A), chitin (Ch), chitosan (Cs), and β-1,6-glucan (H). Three subtypes of α-1,3-glucan (A^a^, A^b^, and A^c^) are indicated by superscripts. Each cross peak represents the correlation between two unique carbons within a molecule, e.g., A^c^1-4 denotes carbon-1 to carbon-4 correlation in type-c α-1,3-glucan. (**b**) Simplified structural representation of rigid polysaccharides, annotated with NMR abbreviations and labeled key carbon positions. (**c**) Molar composition of rigid polysaccharides is estimated from resolved cross-peak volumes in 2D ^13^C CORD spectra. (**d**) Overlay of 2D ^13^C refocused J-INADEQUATE spectra obtained via cross polarization (CP) for rigid molecules of C. *neoformans* and *C. gattii* samples reveal the unique absence of type-c α-1,3-glucan signals (A^c^1 and A^c^2) in *C. gattii cda1*Δ*2*Δ*3*Δ mutant. (**e**) Bright field (BF) and fluorescein-conjugated wheat germ agglutinin (FITC_WGA) images of wild-type strains and their corresponding *cda1*Δ*2*Δ*3*Δ mutants. All strains were grown in YNB (pH 5) for 48 hours and labeled with FITC_WGA to detect chitooligomers. Compared with their wild_type counterparts, *cda1*Δ*2*Δ*3*Δ mutants of both KN99 and R265 exhibit markedly increased WGA binding throughout the cell wall, consistent with enhanced exposure of chitin oligomers in the absence of chitosan.

These spectra further showed that α-1,3-glucan is the dominant rigid polysaccharide in the cell walls of *C. neoformans* KN99 and *C. gattii* R265 (**Fig. 2a, b** and **Supplementary Fig. 1**). Quantitative analysis of cross-peak volumes indicated that α-1,3-glucans account for approximately 71-76% of the rigid polysaccharide fraction in *C. neoformans* and 40-47% in *C. gattii*, confirming their role as the principal structural component of the cryptococcal cell wall (**Fig. 2c** and **Supplementary Table 2**). Chitin and β-1,6-glucan were also detected, accounting for 3-13% and 7-25% of the rigid polysaccharides, respectively, with both present at higher relative abundance in *C. gattii* compared to *C. neoformans*.

Within the rigid α-1,3-glucan pool, three magnetically distinct subtypes (designated A^a^, A^b^, and A^c^) were resolved in both species (**Fig. 2a, d** and **Supplementary Fig. 2**)^35^. These spectroscopic subtypes likely arise from differences in local conformation and interactions with surrounding wall components. In the wild-type cells of both *C. neoformans* and *C. gattii*, type-a α-1,3-glucan represented the most abundant form, whereas types-b and -c were present at lower levels (**Fig. 2c**). The ^13^C chemical shifts of C3 in types a and b (85-86 ppm ppm) align with those reported for aggregated structures such as bundles and sheets, whereas the 82 ppm C3 chemical shift of type-c suggests a non-aggregated single-chain structure^37^. This is consistent with the observation that type-c α-1,3-glucan spans both rigid and mobile domains and acts as a structural bridge between the rigid glucan/chitin/chitosan core and the surrounding soft cell wall matrix (**Supplementary Fig. 3)**.

Chitosan deficiency markedly altered the composition of rigid polysaccharides. In both mutants, we consistently observed a two-fold increase in chitin, from 3% to 7% in *C. neoformans* and from 6% to 13% in *C. gattii* (**Fig. 2d**). Strikingly, the combined amount of chitin and chitosan remained unchanged, totaling 7-8% in both wild-type and mutant strains of *C. neoformans* and 13% in *C. gattii*. These results indicate that deletion of the chitin deacetylase genes affects only the deacetylation of chitin to chitosan, without altering the overall level of chitin-derived polysaccharides.

The other polysaccharides responded differently in the two species. In *C. neoformans*, β-1,6-glucan decreased approximately two-fold, from 15% to 7%, whereas in *C. gattii* its abundance remained largely unchanged in the absence of chitosan (**Fig. 2c**). In addition, *C. neoformans* showed a moderate increase in total α-1,3-glucan, while *C. gattii* exhibited a decrease in α-1,3-glucan, including a complete loss of type-c α-1,3-glucan in the mutant (**Fig. 2a, d**). These changes suggest that chitosan depletion reorganizes the rigid cell wall network differently in the two species and that type-c α-1,3-glucan may rely on chitosan for structural stabilization in *C. gattii*.

The chitosan loss also altered the exposure of chitin-derived PAMPs. Mutants of both species displayed a marked increase in fluorescein-conjugated wheat germ agglutinin (FITC□WGA) binding, indicating increased chitooligomers content or accessibility (**Fig. 2e**). In wild-type cells, WGA staining was largely restricted to bud scars or budding sites, whereas in *cda1*Δ*2*Δ*3*Δ mutants WGA bound broadly across the cell surface in addition to the bud site. These results suggest either elevated levels of chitooligomers or increased accessibility of chitooligomer binding sites in the absence of chitosan.

### Chitosan deficiency does not impair capsule formation but alters its polysaccharide structure

We then examined whether chitosan deficiency affects the global level capsule synthesis and attachment to the cell surface. India ink staining revealed that both wild-type and chitosan-deficient cells of both *Cryptococcus* species produced prominent capsules, visible as exclusion zones surrounding the cells (**Fig. 3a**). This was further confirmed by immunofluorescence labeling with a capsule-specific monoclonal antibody 18B7 (**Fig. 3b**). Comparable capsule morphology and antibody labeling were observed in both wild-type and mutant strains of *C. neoformans* and *C. gattii*, indicating the absence of chitosan does not significantly impair capsule synthesis or its attachment to the cell surface.

**Figure 3.**
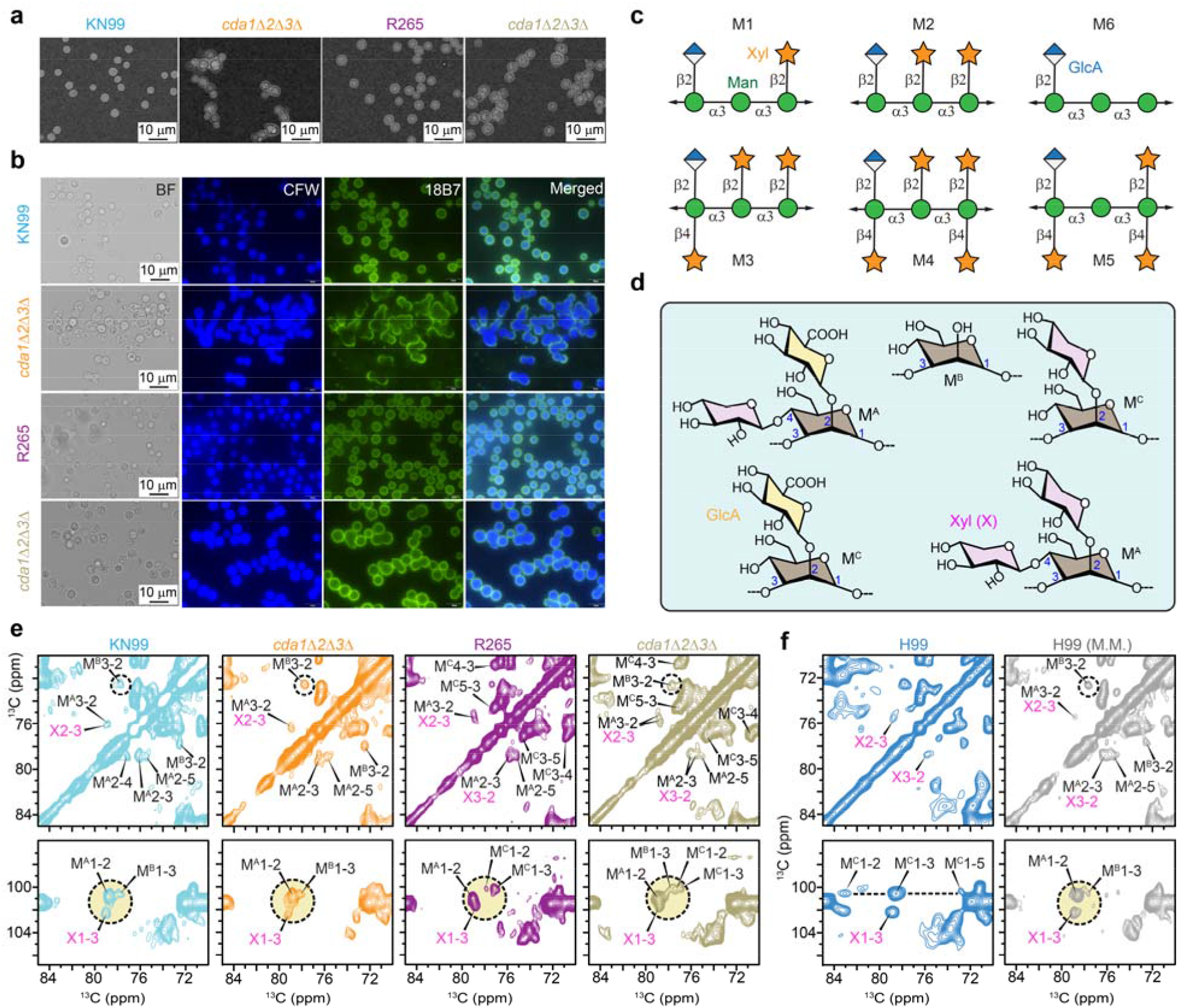
Capsular polysaccharides in different *C. neoformans and C. gattii* strains. Wild-type strains and their corresponding *cda1Δ2Δ3Δ*mutants that were subjected to NMR analysis were stained with either (**a**) India Ink or (**b**) the capsule-specific monoclonal antibody 18B7. For antibody staining, cells were incubated with 18B7 followed by calcofluor white (CFW) and an Alexa Fluor 488-conjugated mouse secondary antibody. Labeled cells were imaged by fluorescence microscopy, and all fluorescent images were acquired using identical exposure settings. Absence of chitosan does not significantly affect capsule production or capsule attachment on the cell surface. (**c**) Representation of six structural motifs of GXM via Symbol Nomenclature for Glycans (SNFG), showing different branching patterns of xylose sidechains. (**d**) Simplified representation of chemical structures of capsular polysaccharides with different linkages of mannose. 1, 2, 3, 4-linked mannose (M^A^); 1, 3-linked mannose (M^B^); and 1, 2, 3-linked mannose (M^C^). (**e**) GXM signals in 2D ^13^C CORD spectra for serotype-A *C. neoformans* KN99 (cyan), its chitosan-deficient mutant (orange), serotype-B *C. gattii* R265 (purple), and its corresponding chitosan-deficient mutant (pale yellow). (**f**) Comparison of GXM signals observed in serotype A *C. neoformans* H99 grown in defined media (blue) and in minimal media (M.M.; grey). Black open circles highlight characteristic signals of type-B mannose. Dashed circles filled with light yellow in the bottom panels show the distinct signals of mannose and xylose. Different types of mannose linkages are observed across all strains.

These observations of capsule formation are consistent with results obtained from solid-state NMR analysis, which provides additional insight into structural variations of capsular polysaccharides. Prior analyses of isolated carbohydrates have shown that the major capsular polysaccharide of *Cryptococcus*, glucuronoxylomannan (GXM), exhibits considerable structural diversity arising from variations in xylose substitution patterns along the mannan backbone, resulting in six distinct structural motifs (M1-M6 in **Fig. 3c**)^38-41^. These structural variations give rise to distinct mannose linkages within the GXM backbone, including 1,2,3,4-linked mannose (labeled as M^A^), 1,3-linked mannose (M^B^), and 1,2,3-linked mannose (M^C^) (**Fig. 3d**). The corresponding signals of these mannose residues in GXM backbone as well as the associated xylose sidechains were observed in the rigid fraction of both *C. neoformans* and *C. gattii*, indicating that certain capsular components can be rigidified through interactions with the cell wall (**Fig. 3e** and **Supplementary Fig. 4**).

However, distinct mannose linkage patterns were observed between the two species. In *C. neoformans* KN99 (serotype A), two forms of mannoses were detected: Mn^A^ corresponds to an α-1,3-linked mannose substituted with glucuronic acid and/or xylose through β-1,2 and β-1,4 linkages, whereas Mn^B^ represents an α-1,3-linked mannose lacking both glucuronic acid and xylose substitutions. The presence of these two mannose environments is consistent with the M5 GXM structural motif. In contrast, *C. gattii* R265 (serotype B) exhibited two different mannose linkages, Mn^A^ and Mn^C^, where the latter corresponds to an α-1,3-linked mannose bearing a single glucuronic acid or xylose branch. Therefore, the GXM of *C. gattii* is more extensively substituted with xylose, consistent with the M4 motif, highlighting species-specific capsule architecture.

Analysis of the chitosan-deficient mutants revealed that the overall GXM linkage pattern in the KN99 *cda1*Δ*2*Δ*3*Δ mutant remained similar to that of the wildtype, with both Mn^A^ and Mn^B^ present. However, in the R265 *cda1*Δ*2*Δ*3*Δ mutant, an additional Mn^B^ signal corresponding to an α-1,3-linked mannose lacking glucuronic acid and xylose substitutions was detected, alongside the Mn^A^ and Mn^C^ forms originally present in the background strain. The appearance of this mannose environment suggests that cell wall reorganization associated with the absence of chitosan have influenced GXM substitution patterns in *C. gattii*, likely as an adaptive response.

We also found that growth conditions influence capsular architecture, with GXM exhibiting distinct linkage patterns even within the same species. In *C. neoformans* H99 grown in defined YNB medium, NMR spectra revealed a single dominant mannose environment (Mn^C^) consistent with the M2 structural motif (**Fig. 3f**). In contrast, when H99 was grown in low-nutrient minimal medium, Mn^A^ and Mn^B^ were detected, resembling the capsular architecture observed in KN99. These results indicate that capsular GXM structure is flexibly remodeled depending not only on species or serotype but also on strain background and environmental growth conditions.

### Depletion of chitosan remodels the mobile cell wall matrix

Although chitosan is part of the rigid cell wall fraction, its loss markedly altered the composition of the mobile cell wall domain. In the mobile fractions of both *C. neoformans* and *C. gattii*, β-1,6-glucan was the predominant component, accompanied by weaker contributions from its β-1,3-glucan sidechains, as well as type-c α-1,3-glucan and mannoproteins (**Fig. 4a, b**). Chitosan deficiency led to a pronounced reduction in β-1,6-glucan backbone in *C. neoformans*, decreasing from 79% to 53% in the mobile fraction, whereas the reduction was more moderate in *C. gattii* (**Fig. 4c** and **Supplementary Table 3**), consistent with earlier observations in the rigid fraction (**Fig. 1c**). In addition, β-1,3-linked side chains of the β-glucan matrix were completely depleted in the chitosan mutant of *C. neoformans*, whereas they doubled in the mutant strain of *C. gattii*, increasing from 4% to 8% (**Fig. 4a, c**). These results suggest that β-glucans are more functionally coupled to chitosan-dependent cell wall organization in *C. neoformans*, whereas *C. gattii* appears to maintain β-glucan composition more robustly following chitosan loss.

**Figure 4.**
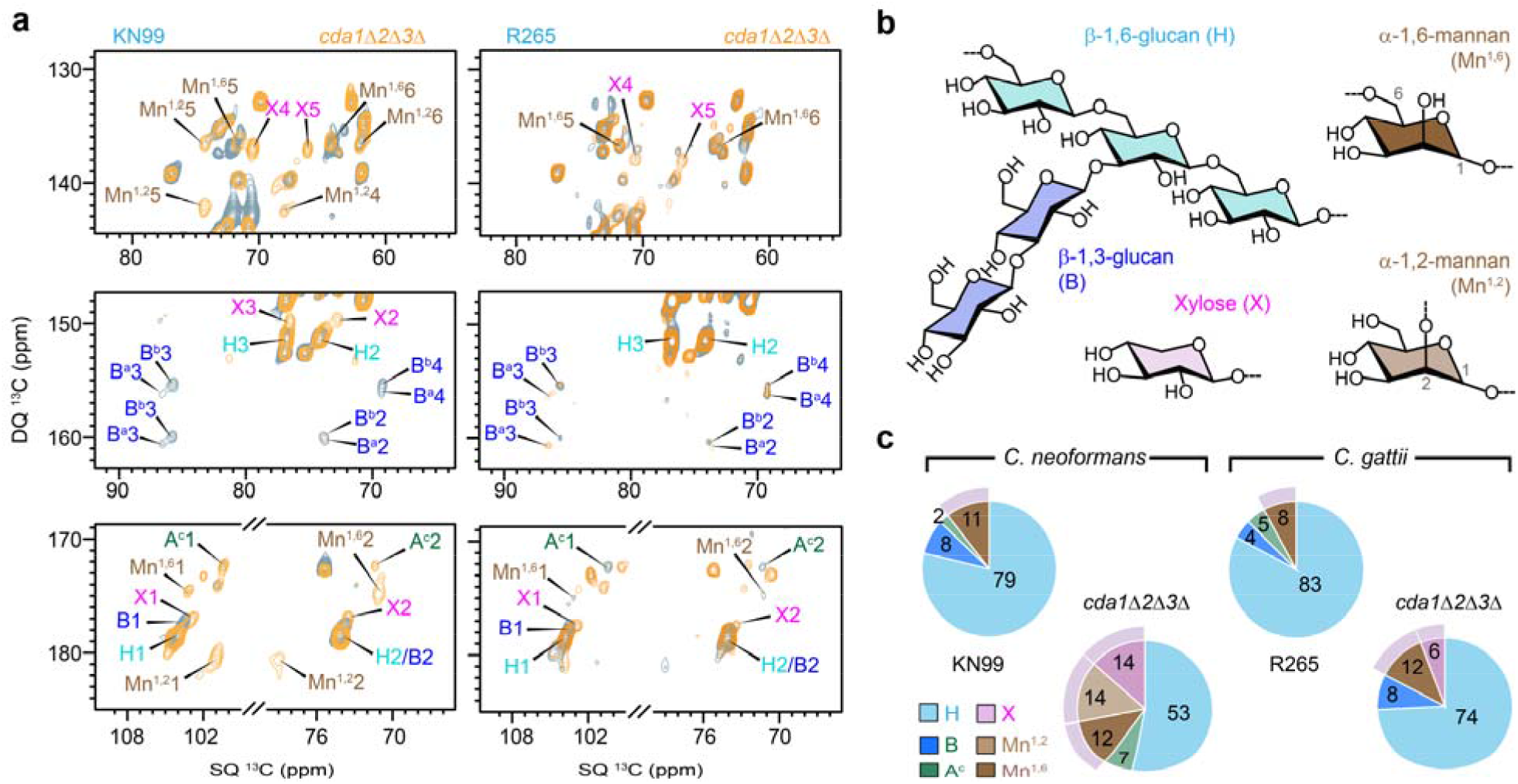
Mobile polysaccharides in wildtype and mutant strains of *C. neoformans* and *C. gattii*. (**a**) Overlay of 2D ^13^C refocused J-INADEQUATE spectra obtained via direct polarization (DP) and short recycle delays for selectively detecting mobile molecules in KN99 (left, blue) and its *cda1*Δ*2*Δ*3*Δ mutant (left, yellow), as well as in R265 (right, blue) and its *cda1*Δ*2*Δ*3*Δ mutant (left, yellow). (**b**) Representative structures of β-glucans and mannoproteins (α-1,2-mannan, α-1,6-mannan, and xylose) in *Cryptococcus*. (**c**) Molar composition of mobile polysaccharides estimated from well-resolved peaks in 2D ^13^C DP refocused J-INADEQUATE spectra.

Conversely, type-c α-1,3-glucan content increased markedly in the mutant cells of *C. neoformans*, but was depleted in the mutant of *C. gattii* (**Fig. c**). These reciprocal changes indicate distinct compensatory remodeling strategies in the two species following disruption of chitosan. In *C. neoformans*, enhanced α-1,3-glucan mobility may partially compensate for the loss of β-1,3-glucan, whereas *C. gattii* preserves β-glucan content while eliminating mobile α-1,3-glucan.

Chitosan deficiency also induced pronounced changes in mannoprotein composition. Cryptococcal mannoproteins consist primarily of α-1,2- and α-1,6-linked mannoses with xylose substitutions^42-44^. In *C. neoformans*, the overall mannoprotein content increased in the mutant, with new signals corresponding to α-1,2-linked mannose detected (**Fig. 4a, c**). Differently, the mutant cells of *C. gattii* showed an increase in mannoproteins mainly characterized by enrichment of α-1,6-linked mannose, rising from 8% to 12%, while α-1,2-linked mannose signals remained minimal (<1%). Additionally, xylose signals were detected in the mobile fractions of both chitosan-deficient mutants, which may originate from either GXM or xylose-containing mannoproteins (**Fig. 4a, b**). Notably, the chemical shifts of xylose in the mobile domain differed from those observed in the rigid domain (**Supplementary Table 1**), indicating that these residues have distinct chemical and structural environments.

### *cda1*Δ*2*Δ*3*Δ mutant showed increased chitin rigidity and reduced water accessibility

While the cell wall architecture of *C. neoformans* has been characterized in a recent solid-state NMR study, the organization of the *C. gattii* cell wall remains poorly understood^35^. Therefore, we performed a 2D proton-assisted recoupling (PAR) experiment on *C. gattii* R265 to probe intermolecular spatial proximities among cell wall polysaccharides (**Fig. 5a**)^45,46^. Notably, signals from chitosan and capsular polysaccharides are absent in the PAR spectrum (asterisks in **Fig. 5a**), likely due to their semi-disordered nature. Consequently, the detected correlations arise predominantly from interactions among crystalline chitin microfibrils and rigid α-glucans.

**Figure 5.**
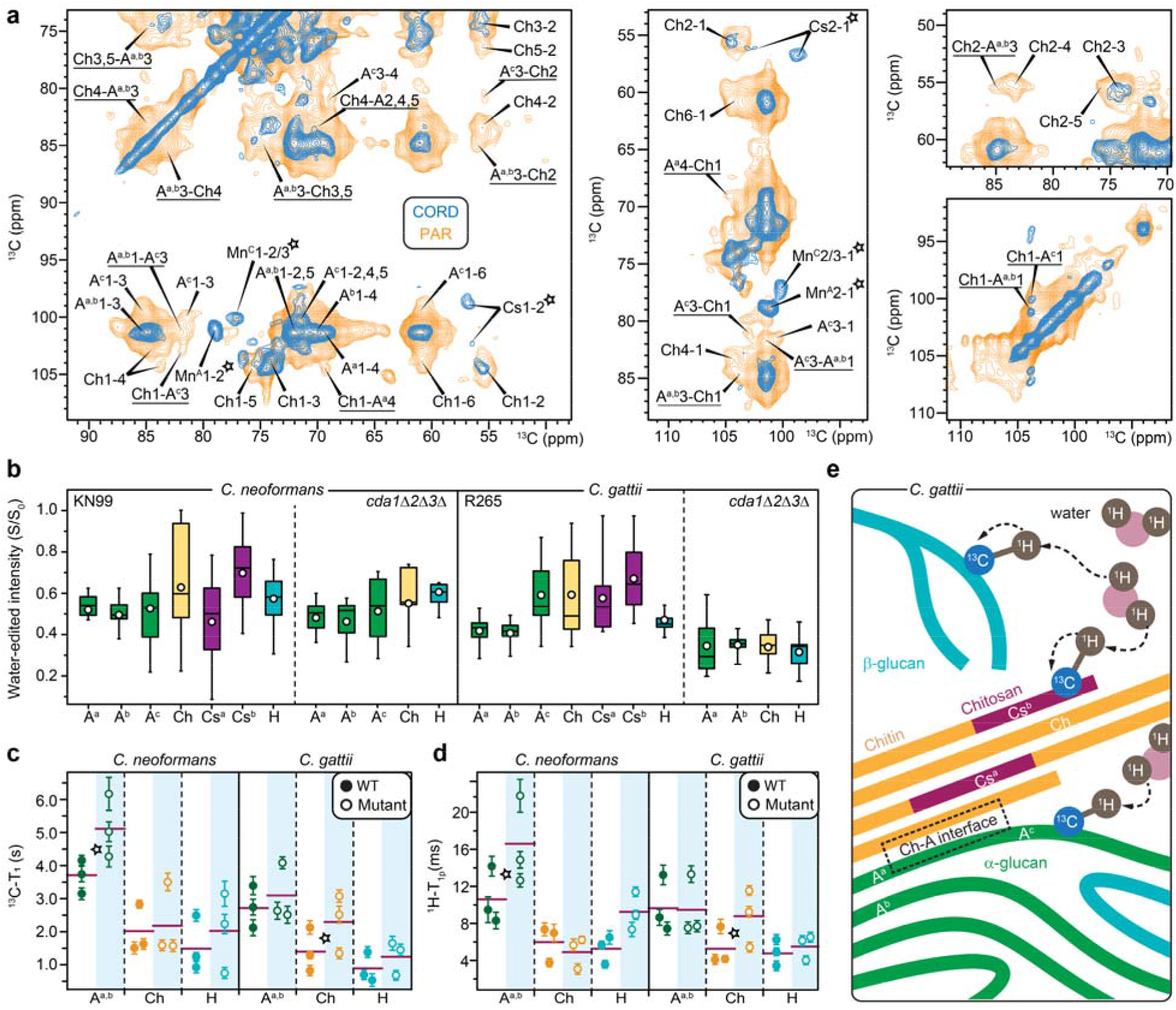
Polysaccharide packing, water accessibility, and dynamics in *C. neoformans* and *C. gattii*. (**a**) Intermolecular interactions of polysaccharides probed by 2D ^13^C-^13^C correlation spectrum. The 53-ms CORD spectrum (blue) detects intramolecular cross peaks among carbon sites within each molecule, while the 15-ms PAR spectrum (light orange) detects additional, long-range intermolecular cross peaks happening between different molecules. Underlines annotate intermolecular cross peaks uniquely observed in PAR. Open asterisks annotate signals from chitosan and capsular polysaccharides that were absent in PAR spectrum. (**b**) Intensity ratios (S/S_0_) between water-edited peak spectra (S) and control spectra (S_0_) showing the extent of water association for α-1,3-glucan (A), chitin (Ch), chitosan (Cs), and β-1,6-glucan (H), with the superscripts indicating the respective subtypes. Open circles: average; horizontal lines: median. Data size: A^a^ (n=16) in all four samples; A^b^, Ch, H (n=9 each) in all; A^c^ (n=9) in all except R265 *cda1*Δ*2*Δ*3*Δ mutant; Cs^a^, and Cs^b^ (n=9 each) in KN99 and R265 only. (**c**) ^13^C-T_1_ relaxation time constants of rigid cell wall polysaccharides. (**d**) 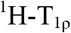 relaxation time constants of rigid polysaccharides. For both panels b and c, error bars are s.d. of the fit parameters and horizontal lines represent the average. (**e**) Concepts of polymer organization in wildtype *C. gattii* restrained by profiles of interactions, hydration, and dynamics. Dashed arrows illustrate ^1^H polarization transfer from water to polysaccharides. Structural polymorphs, such as Cs^a^ and Cs^b^ for chitosan and A^a^, A^b^, and A^c^ for α-glucans are labeled. This is not a cell wall structural model.

The observed correlations arise primarily from intermolecular cross peaks between chitin and α-1,3-glucans. Although signals corresponding to type-c α-1,3-glucan (A^c^) are relatively weak, this polymorph shows clear cross peaks with chitin, including A^c^3-Ch2, Ch1-A^c^3, A^c^3-Ch1, and Ch1-A^c^1. These correlations indicate that type-c α-1,3-glucan is spatially associated with chitin microfibrils. Such contacts may explain why this structural form, which exists as single strands, can span both rigid and mobile domains, with the chitin-contacting portion becoming rigidified. In addition, type-c α-1,3-glucan exhibits cross peaks with the other two α-1,3-glucan polymorphs, such as A^a,b^1-A^c^3 and A^c^3-A^a,b^1, indicating that all three α-1,3-glucan forms are spatially proximal within the rigid domain.

Signals from type-a and type-b α-1,3-glucans largely overlap in the spectra, resulting in numerous observed cross peaks between these forms and chitin, including Ch4-A^a,b^3, A ^a,b^3-Ch4, A ^a,b^3-Ch3,5, A ^a,b^3-Ch1, and Ch2-A^a,b^3. This ambiguity can be partially resolved by examining the C4 carbon sites, where A^b^1-4 can be distinguished from A^a^1-4. Interestingly, cross peaks Ch1-A^a^4 and A^a^4-Ch1 are observed, whereas the corresponding correlations for the A^b^ form are absent. These observations suggest that type-a α-1,3-glucan is more closely packed with chitin, whereas type-b α-1,3-glucan is positioned further away. Given that both A^a^ and A^b^ exhibit chemical shifts consistent with aggregated α-glucan structures, these results indicate that only the A^a^ polymorph forms domains that are directly proximal to chitin microfibrils.

Water accessiblity of rigid polysaccharides was assessed using water-edited ^1^H polarization transfer experiments, in which ^1^H polarization is transferred from bound water to nearby polysaccharides (**Fig. 5b** and **Supplementary Fig. 5**) ^47-49^. The S/S_0_ intensity ratios between water-edited (S) and control (S_0_) spectra report site-specific water accessibility. Among all rigid polysaccharides, type-a and type-b α-1,3-glucans were the least hydrated components in both KN99 and R265 cell walls, with average S/S_0_ values of 0.49-0.52 in KN99, and 0.41-0.42 in R265 (**Fig. 5b** and **Supplementary Table 4**). In KN99, type-c α-1,3-glucan displayed hydration levels comparable to those of type-a and type-b, whereas in R265 it showed noticeably higher hydration. This difference suggests that type-c α-1,3-glucan is better integrated into the rigid α-glucan network in KN99, whereas in *C. gattii* it is loosely associated and therefore more hydrated.

The limited water accessibility of most α-1,3-glucans is consistent with their restricted mobility, as revealed by NMR relaxation measurements (**Fig. 5c, d**; **Supplementary Fig. 6, 7**). Type-a/b α-1,3-glucan exhibited the longest ^13^C-T_1_ and 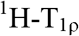 relaxation times in both strains, with average values of 2.7-3.7 s and 9.7-10.6 ms, respectively (**Fig. 5c, d** and **Supplementary Table 5**). These values are substantially greater than those measured for chitin and β-1,6-glucan, indicating that type-a/b α-1,3-glucans undergo highly restricted molecular motions on nanosecond and microsecond timescales. Together with their limited hydration and characteristic C3 chemical shifts, these results support a model in which type-a/b α-1,3-glucans assemble into densely packed supramolecular complexes that largely exclude bulk water and form a highly rigid structural core within the cell wall.

In contrast, type-b chitosan (Cs^b^) was the most hydrated rigid polysaccharide in both KN99 and R265, with average S/S_0_ values of 0.69 and 0.67, respectively (**Fig. 5b** and **Supplementary Table 4**), suggesting that this form contributes to maintaining water accessibility within structural domains where chitin and chitosan coexist. Type-a chitosan (Cs^a^), however, exhibited lower hydration levels, consistent with a more embedded position within the chitin-chitosan fibril/domain.

Chitin exhibited intermediate dyanmics, with average ^13^C-T_1_ of 1.4-2.0 s and averge 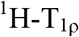 of 5.3-6.0 ms, indicating more restricted motion than β-1,6-glucan but greater flexibility than α-1,3-glucans (**Fig. 5d**,**e** and **Supplementary Table 5**). Although chitin forms microfibrils primarily through antiparallel chain packing, it is not the most rigid or least hydrated polysaccharide in the cryptococcal cell wall. This behavior likely reflects extensive deacetylation of chitin to chitosan, estimated to exceed 50% based on molar composition (**Fig. 2c**), which increases the flexibility of the chitin microfibrillar network.

Finally, *C. neoformans* and *C. gattii* exhibited markedly different responses to chitosan loss. In *C. neoforman*s, hydration levels of rigid polysaccharides in the chitosan-deficient mutant decreased only slightly relative to wild-type KN99. In contrast, *C. gattii* displayed a pronounced reduction in hydration across all rigid polysaccharides, including α-1,3-glucans, β-1,6-glucan, and chitin (**Fig. 5b** and **Supplementary Table 4**). These results indicate that chitosan plays a substantially larger role in maintaining hydration of the rigid cell wall matrix in *C. gattii* than in *C. neoformans*. Notably, upon chitosan loss, *C. neoformans* reinforces the cell wall by increasing α-1,3-glucan rigidity, whereas *C. gattii* forms more rigid chitin microfibrils, as indicated by increased ^13^C-T_1_ and 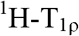 relaxation times in the mutants relative to respective wild-type strains (**Fig. 5c, d**).

These structural data reveal a distinctive organization of cell wall polysaccharides in *C. gattii*, in which chitosan fulfills two complementary roles: the type-b allomorph associates with water and contributes to hydration of the rigid matrix, whereas the type-a form maintains flexibility within the chitin-chitosan fibrillar core (**Fig. 5e**). Loss of these chitosan elements therefore leads to pronounced dehydration of the cell wall. In addition, chitin is extensively associated with type-a and type-c α-1,3-glucans. The type-a form participates in highly aggregated α-glucan domains together with the less accessible type-b polymorph, whereas the type-c form adopts a single-strand structure that bridges the rigid and mobile domains. This more exposed configuration results in higher hydration and additional associations with β-glucans. Notably, β-glucan content is reduced by half in *C. neoformans* upon chitosan loss but remains largely unaffected in *C. gattii*, further highlighting species-specific differences in cell wall remodeling.

## DISCUSSION

This study reveals fundamental differences in the supramolecular organization of the cell walls of *C. neoformans* and *C. gattii*. Using solid-state NMR spectroscopy on intact fungal cells, we demonstrate that although both species rely on α-1,3-glucans as the principal rigid scaffold, the relative contributions of chitin, chitosan, and β-glucans differ substantially between them (**Fig. 2c**). In particular, *C. gattii* exhibits a more complex multicomponent architecture in which chitin, chitosan, and β-glucan play greater roles in maintaining cell wall integrity. These results reveal that closely related cryptococcal pathogens employ distinct structural strategies to assemble and maintain their cell walls.

One particularly notable difference between the two species concerns the behavior of type-c α-1,3-glucan. In a recent study of *C. neoformans*, this polymorph was found to bridge the rigid and mobile cell wall domains^35^. Our data show that this bridging role is conserved in *C. gattii*, where type-c α-1,3-glucan exhibits higher hydration than the aggregated α-glucan forms (**Fig. 5b**). However, in the KN99 strain of *C. neoformans*, type-c α-1,3-glucan displays hydration levels similar to those of the other α-glucan polymorphs, indicating tighter integration into the rigid network (**Fig. 5b**). These observations highlight the structural plasticity of the cryptococcal cell wall and suggest that α-glucan polymorphism enables different architectural configurations across strains and species.

Chitosan is known as a more flexible and soluble polymer than chitin and therefore makes the cell wall more flexible and adaptive to environmental stresses^18,22,23,50^. Our data further demonstrate that chitosan plays a central role in organizing the cryptococcal cell wall matrix.

Two chitosan subtypes were resolved spectroscopically, which differ in hydration and structural integration (**Figs. 2a** and **5b**). Type-b chitosan is consistently more hydrated and appears to maintain water accessibility within chitin-chitosan fibrillar domains, whereas type-a chitosan is more deeply embedded within the fibrillar core. These findings indicate that chitosan not only contributes to mechanical stability but also modulates the hydration properties of the rigid cell wall matrix. Consistent with this role, deletion of the three chitin deacetylase genes markedly reduced hydration of rigid polysaccharides in *C. gattii*, whereas the corresponding changes in *C. neoformans* were comparatively modest, relying on its β-glucan matrix to maintain hydration (**Fig. 5b**).

Chitosan depletion triggers extensive yet species-specific remodeling of the cell wall. In *C. neoformans*, loss of chitosan leads to increased α-1,3-glucan and chitin in the rigid scaffold, accompanied by a reduction in β-1,6-glucan and complete loss of β-1,3-glucan from the mobile domain (**Figs. 2c** and **4c**). In contrast, *C. gattii* responds through a distinct remodeling strategy characterized by loss of type-c α-1,3-glucan, increased β-1,3-glucan, and extensive reorganization of mannoproteins. These divergent responses indicate that the two species rely on different structural networks to maintain cell wall integrity when chitosan biosynthesis is disrupted.

Chitosan depletion also influences extracellular polysaccharides associated with the cell surface. Although capsule production itself was not impaired, solid-state NMR analysis revealed species-specific changes in the structure of capsular GXM (**Fig. 3e**). The structure of cryptococcal GXM is highly variable and depends on both strain background and environmental growth conditions (**Fig. 3e**). Early compositional analyses of GXM isolated from culture supernatants identified distinct molar ratios of xylose, mannose, and glucuronic acid, approximately 1:3:1, 2:3:1, 3:3:1, and 4:3:1 for serotypes D, A, B, and C, respectively, corresponding primarily to motifs M1-M4^38,51^. Subsequent studies demonstrated that individual strains produce heterogeneous mixtures of motifs; for example, serotype A strains H99 and KN99 contain multiple motifs within their exopolysaccharide fractions, with H99 composed predominantly M2 (60%) alongside M3 (25%) and M4 (15%), and KN99 containing M1 (15%), M2 (50%), and M4 (35%)^40^. In contrast to these analyses of shed exopolysaccharides, our examination of rigid capsular polysaccharides tightly associated with the cell wall reveals a distinct motif distribution: motifs M2 and M5 in the serotype A strains H99 and KN99, respectively, and motif M4 predominating in the serotype B strain R265 (**Fig. 3c-f**). These findings indicate that the structure of cell wall-associated GXM can differ from that of released capsule material and may be influenced by the underlying cell wall architecture^52^.

In both species, chitosan depletion also led to a pronounced increase in mannoproteins (**Fig. 4a, c**), a key immunogenic component located in the mobile domain of the cryptococcal cell wall^53^. In *Saccharomyces* and *Candida* species, mannoproteins are covalently and non-covalently associated with surface glucans, whereas in *Cryptococcus*, their positioning within the cell wall is highly dynamic^54-57^. Immunoelectron microscopy has revealed an asymmetric distribution of mannoproteins, supporting a model in which mannoproteins migrate slowly through the dense inner cell wall layer but diffuse more rapidly and are released upon reaching the less compact outer layer^58^. Importantly, engagement of cryptococcal mannoproteins with mannose receptors on dendritic cells promotes T-cell activation and confers protective immunity against *C. neoformans*, leading to their proposal as vaccine candidates for cryptococcosis^59-61^. Consistent with this concept, our solid-state NMR analyses reveal a pronounced increase in mannoproteins in chitosan-deficient mutants of both *C. neoformans* and *C. gattii* (**Fig. 4a, c**), suggesting that enhanced mannoprotein exposure and release may underline the host-protective immune responses observed in these avirulent strains.

Together, these findings support a model in which the cryptococcal cell wall is organized as a hierarchical network of interacting polysaccharides. Aggregated α-1,3-glucans form a rigid structural core closely associated with chitin microfibrils, while chitosan regulates hydration and flexibility within the chitin-glucan interface (**Fig. 5e**). Surrounding this scaffold is a more dynamic matrix of β-glucans and mannoproteins that can be remodeled in response to perturbations in cell wall composition. Within this framework, the two species adopt distinct architectural strategies: *C. neoformans* primarily relies on an α-1,3-glucan-dominated scaffold, whereas *C. gattii* depends on a more multicomponent network involving chitin, chitosan, and β-glucans. Chitosan depletion triggers species-specific remodeling of the cell wall matrix, reshaping polymer interactions and hydration properties and potentially altering the exposure of PAMP including chitin-derived oligomers and mannoproteins^62^. The species-specific structural strategies uncovered here may help explain differences in pathogenicity, drug susceptibility, and immune recognition between cryptococcal pathogens and emphasize the importance of understanding cell-wall architecture and remodeling when developing antifungal therapies and vaccines^63^.

## MATERIALS AND METHODS

### Preparation of uniformly ^13^C,^15^N-labeled fungal material

*C. neoformans* KN99, and its *cda1*Δ*2*Δ*3*Δ mutant, and *C. gattii* R265, and its *cda1*Δ*2*Δ*3*Δ mutant were cultured in ^13^C, ^15^N-enriched growth media. The growth medium consisted of 0.67% Yeast Nitrogen Base (YNB), 2.0% ^13^C-glucose (Cambridge Isotope Laboratories, CLM-1396-PK), and 1.5% ^15^N-ammonium sulfate (Cambridge Isotope Laboratories, NLM-713-PK), with the pH adjusted to 5.5 using 1M HEPES buffer. Cells were cultured in 100 mL liquid medium in 250-mL Erlenmeyer flasks and incubated at 30°C with shaking at 300 rpm (1,364 × g). Fungal biomass was harvested by centrifugation at 4,000 rpm (3,700 × g) for 5 min at 4°C and the pellet was washed four times using nano-purified water followed by the centrifugation procedures. For solid-state NMR characterization, 35-45 mg of natively hydrated whole-cell material was packed into a 3.2 mm magic-angle spinning (MAS) rotor (Cortecnet, HZ16916) for each sample.

### Transmission electron microscopy

For ultrastructural analyses, cells from the culture that was used for NMR analysis were collected by centrifugation at 3600xG for 10 min. Pelleted cells were fixed in 2% paraformaldehyde/2.5% glutaraldehyde (Polysciences Inc., Warrington, PA) in 100 mM sodium cacodylate buffer, pH 7.2 for 2 h ice and then overnight at 4°C. Samples were washed in sodium cacodylate buffer and postfixed in 1% osmium tetroxide (Polysciences Inc.) for 1 h at room temperature. Samples were then rinsed extensively in dH_2_0 prior to en bloc staining with 1% aqueous uranyl acetate (Ted Pella Inc., Redding, CA) for 1 h. Following several rinses in dH_2_0, samples were dehydrated in a graded series of ethanol and embedded in Eponate 12 resin (Ted Pella Inc.). Sections of 95 nm were cut with a Leica Ultracut UC7 ultramicrotome (Leica Microsystems Inc., Bannockburn, IL), stained with uranyl acetate and lead citrate, and viewed on a JEOL2100Plus transmission electron microscope (JEOL USA Inc., Peabody, MA) equipped with an AMT 8-megapixel digital camera and AMT Image Capture Engine V602 software (Advanced Microscopy Techniques, Woburn, MA).

### Cell wall and capsule staining

A portion of the culture used for NMR analysis was processed for cell wall and capsule staining. Cells were harvested by centrifugation at 3500 × g for 8 min at 4□°C and fixed in 4% paraformaldehyde for 30 minutes on ice. Fixed cells were washed twice with phosphate-buffered saline (PBS) and resuspended to a final density of OD□□□ = 1.0. For each staining condition, 100□µL of the cell suspension was used.

Prior to staining, cells were blocked with PBS containing 2% bovine serum albumin (BSA; Sigma-Aldrich, A7906) for 30 min at 30□°C with gentle nutation. For cell wall staining, Calcofluor White (CFW) and Alexa Fluor 488–conjugated wheat germ agglutinin (WGA; Invitrogen™, W11261) were used at final concentrations of 5□µg/mL and 100□µg/mL, respectively. Following staining, cells were washed three times with PBS and spotted onto microscope slides for imaging.

For capsule immunofluorescence staining, cells were incubated with monoclonal antibody 18B7 at a final concentration of 1□µg/mL for 1□h at 4□°C. Cells were then washed twice with PBS containing 2% BSA and incubated with a goat anti-mouse FITC–conjugated secondary antibody (Santa Cruz Biotechnology, SC2010) at 4□µg/mL for 1□h at 4□°C. After incubation, cells were washed twice with PBS containing 2% BSA and prepared for imaging.

Fluorescence microscopy was performed using a 63× objective on an Olympus microscope. WGA fluorescence was detected using GFP filter settings, and CFW fluorescence was detected using 4′,6-diamidino-2-phenylindole (DAPI) filter settings on a Zeiss Axio Imager M2 microscope equipped with a Hamamatsu Flash4.0 CMOS camera.

Capsule staining by India ink was performed by mixing 50□µL of cell suspension from the NMR culture with 50□µL of India ink (STIIN25; American Master Tech Scientific). The mixture was immediately used for microscopy.

### ^13^C solid-state NMR experiments

Most high-resolution 1D and 2D solid-state NMR experiments were conducted at 800 MHz (18.8 T) Bruker Avance Neo spectrometer at Max T. Roger NMR facility of Michigan State University. All ^13^C-detection experiments were performed using a 3.2 mm HCN triple-resonance probe at 15 kHz MAS, with ambient temperatures between 283 K and 298 K. ^13^C chemical shifts were externally referenced by calibrating the N-formyl-L-Met-L-Leu-L-Phe-OH (MLF) methionyl CH_3_ peak to 14.0 ppm, and the resulting spectral reference (sr) value was applied to the spectra of fungal samples collected within the same week. Unless otherwise specified, typical radiofrequency field strengths were 83-100 kHz for ^1^H decoupling, 62.5 kHz for ^1^H hard pulses, and 50-62.5 kHz for ^13^C. Experimental parameters for all NMR spectra are documented in **Supplementary Table 6**.

To facilitate resonance assignment, 2D ^13^C-^13^C correlation experiment was conducted using a 53-ms CORD (combined R2_n_^v^-driven) mixing period, which revealed intramolecular cross-peaks between carbon sites within each molecule (**Supplementary Fig. 1**)^36^. Additionally, 2D ^13^C refocused J-INADEQUATE experiments were conducted using either CP or DP (1.5 s recycle delay) to probe molecular domains with different mobilities (**Supplementary Fig. 3**)^64^. This experiment correlated double-quantum (DQ) chemical shifts with two corresponding single-quantum (SQ) shifts, generating asymmetric spectra that enabled efficient tracking of through-bond carbon-connectivity. Each τ period (out of four) was set to 2.3 ms to maximize the intensities of carbohydrate peaks. The 2D CORD experiment was conducted on all four *Cryptococcus* strains. The assigned chemical shifts for rigid and mobile polysaccharides are documented in **Supplementary Table 1**. Data acquisition was conducted using Topspin 3.5, and spectral analysis was performed in Topspin 4.2.0. Figures were prepared using Adobe Illustrator CS6 (V16.0.0).

Relative molecular composition analysis of carbohydrates was conducted by selecting only well-resolved signals in 2D ^13^C CORD spectra for rigid components and ^13^C DP refocused J-INADEQUATE spectra for mobile molecules (**Supplementary Table 2** and **3**)^64^. Peak volumes were quantified using the integration function in the Bruker Topspin software, with quantification based on the mean of the peak volumes from resolved signals. All spectra used for comparison were collected and processed under the same pulse sequences, acquisition parameters, and processing conditions, with normalization applied based on the number of scans. The relative abundances of different polysaccharides were determined by normalizing the sum of integrals with their respective counts. The standard error for each polysaccharide was calculated by dividing the standard deviation of integrated peak volumes by the total cross-peak count. The overall standard error was computed as the square root of the sum of squared errors for each polysaccharide. The percentage error was determined by normalizing the standard error with the average integrated peak volume and adjusting for each polysaccharide’s relative abundance^65^.

### Solid-state NMR of polymer hydration and dynamics

To probe the water-accessibility of cell wall polymers, 1D and 2D water-edited ^13^C-^13^C correlation experiments^47,49^ were conducted on a Bruker Avance Neo 400 MHz (9.4 T) NMR spectrometer at Michigan State University using a 3.2 mm HCN MAS Bruker probe at 280 K. Briefly, a ^1^H-T_2_ filter (0.6-1.2 ms, strain-dependent) was applied to suppress carbohydrate signals to less than 10% but preserve 80-93% of water magnetization. (**Supplementary Fig. 5**), followed by ^1^H-^1^H mixing to transfer water ^1^H magnetization to hydrated carbohydrates. ^13^C detection was achieved via CP with a 1-ms contact time. For 2D water-edited experiments, a 4 ms ^1^H mixing period and 50 ms DARR mixing were used. Intensity ratios (S/S_0_) between the water-edited (S) and control (S_0_) spectra were obtained to assess water retention of each carbon site (**Supplementary Table 4**).

To probe polymer dynamics, ^13^C-T_1_ relaxation was measured using the Torchia-CP scheme^66^ with a varied z-filter duration ranging from 0.1 μs to 8 s. For ^13^C-detected, 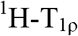 relaxation, a Lee-Goldburg (LG) spinlock sequence combined with LG-CP suppressed ^1^H spin diffusion, enabling site-specific measurements via bonded ^13^C detection^67,68^. For both measurements, peak intensity decay was fitted to a single exponential equation to determine the corresponding relaxation time constants (**Supplementary Figs. 6, 7**and **Supplementary Table 5**). The analysis was conducted using OriginPro 9.

### Solid-state NMR analysis of inter-polysaccharide interactions

Approximately 30 mg of *C. gattii* cells was packed into a 3.2-mm pencil rotor for intermolecular interaction analysis. The 2D ^13^C PAR experiment^45,46^, which is power-demanding, was conducted on a 800 MHz Bruker Avance Neo spectrometer at the National High Magnetic Field Laboratory (Tallahassee, FL, USA). Data acquisition utilized a 3.2-mm HCN triple-resonance probe operating at 15 kHz MAS and 280 K. The PAR spectrum was acquired with a 15 ms recoupling period, during which the ^1^H and ^13^C irradiation frequencies were set at 53 kHz and 50 kHz, respectively.

## DATA AVAILABILITY

All relevant data that support the findings of this study are provided in the article and Supplementary Information. All the original ssNMR data files will be deposited in the Zenodo repository, and the access code and DOI will be provided. Source Data will be provided as a source data file.

## AUTHOR CONTRIBUTIONS

A.A., M.D., L.X., and I.H. conducted solid-state NMR experiments and data analysis. A.A. and R.U. prepared fungal samples. R.U. and D.F. performed imaging analysis. A.A. and R.U. wrote the first draft of the manuscript. J.K.L. and T.W. designed and supervised the project. All authors contributed to the manuscript writing.

## ACKNOWLEDGMENT

This study was primarily supported by National Institutes of Health (NIH) under award number R01AI173270 to T.W and R01AI123407 to J.K.L. A portion of this work was performed at the National High Magnetic Field Laboratory, which is supported by National Science Foundation Cooperative Agreement No. DMR-2128556 and the State of Florida. The electron microscopy was performed on a JEOL2100Plus, funded by a NIH Shared Instrumentation Grant (to Sara E Miller, Director Center for Electron Microscopy & Nanoscale Technology, Duke University, Grant number 1S10OD026776).

