## Supplementary file for "Molecular Architecture of *Cryptococcus* Cell Walls Reveals Species-Specific Chitosan-Dependent Remodeling"

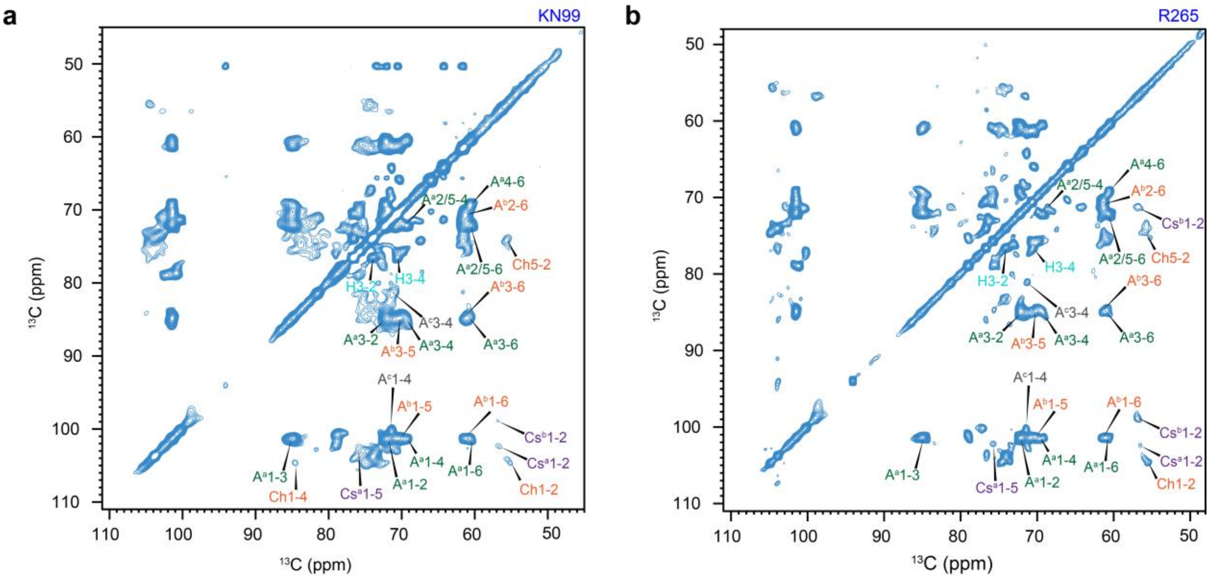


**Supplementary Figure 1. Resonance assignment of rigid glucans in *C. neoformans* and *C. gattii* cell wall.** CP-based 2D $^{13}$C-$^{13}$C correlation spectrum measured with 53 ms CORD mixing for **(a)** KN99, and **(b)** R265.

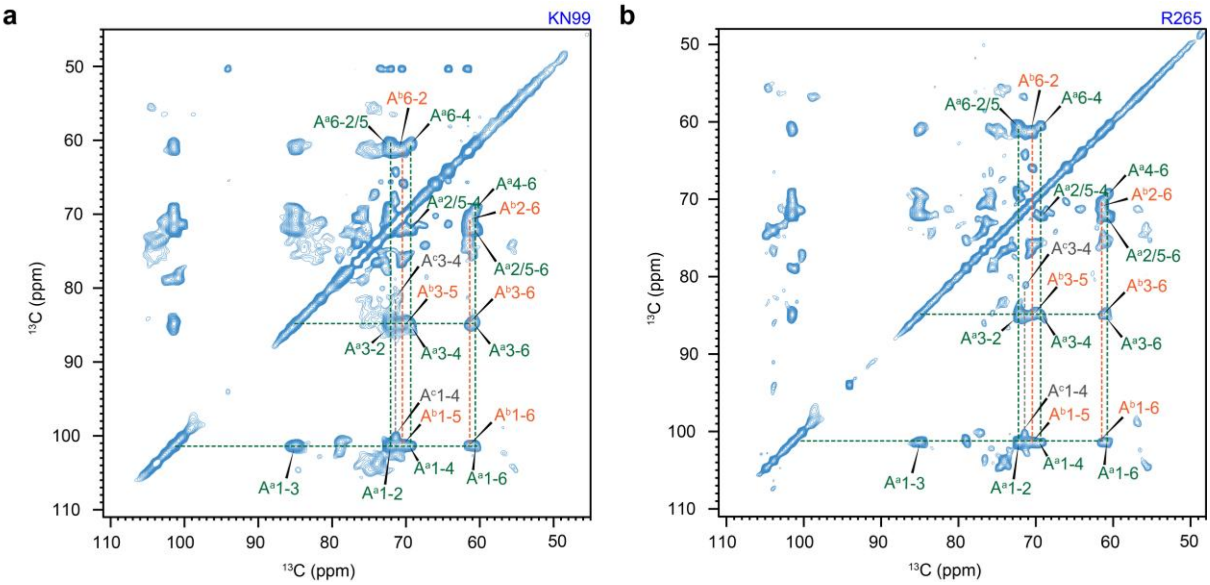


**Supplementary Figure 2. Resonance assignment of α-1,3-glucan in *C. neoformans* and *C. gattii* cell wall.** CP-based 2D $^{13}C$-$^{13}C$ correlation spectrum measured with 53 ms CORD mixing. **(a)** KN99, and **(b)** R265. Each peak is annotated with the abbreviation of the carbohydrate name, the subtype (in superscript), and the carbon number. For instance, $A^c1$ represents the carbon 1 of type-c α-1,3-glucan.

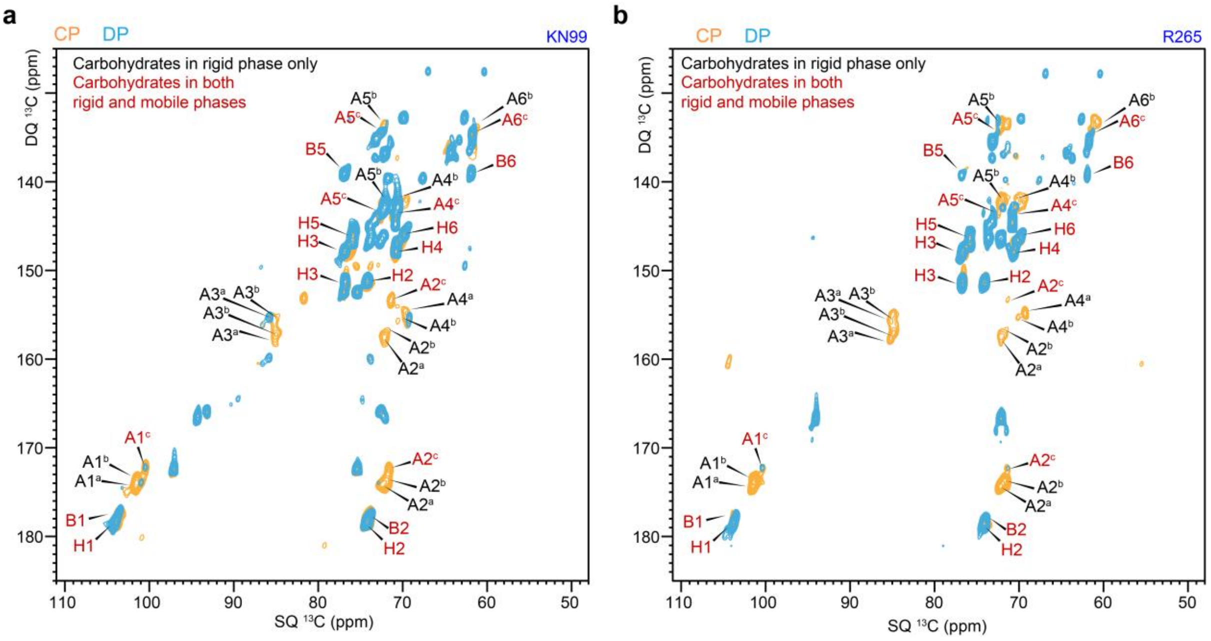


**Supplementary Figure 3. Distribution of α-1,3-glucan, and β-glucan in rigid and mobile domains.** Overlay of 2D refocused J-INADEQUATE spectra measured with CP (orange) and DP (cyan) for **(a)** *C. neoformans* KN99 and **(b)** *C. gattii* R265 samples. The carbohydrates observed only in the CP-based spectra are rigid and are marked in black. The carbohydrates observed in both CP-INADEQUATE and CP-INADEQUATE spectra are marked in red: these carbohydrates have two-modal distribution in rigid and mobile phases. β-1,6-glucan, β-1,3-glucan, and types-c α-1,3-glucan were observed in both domains for both samples.

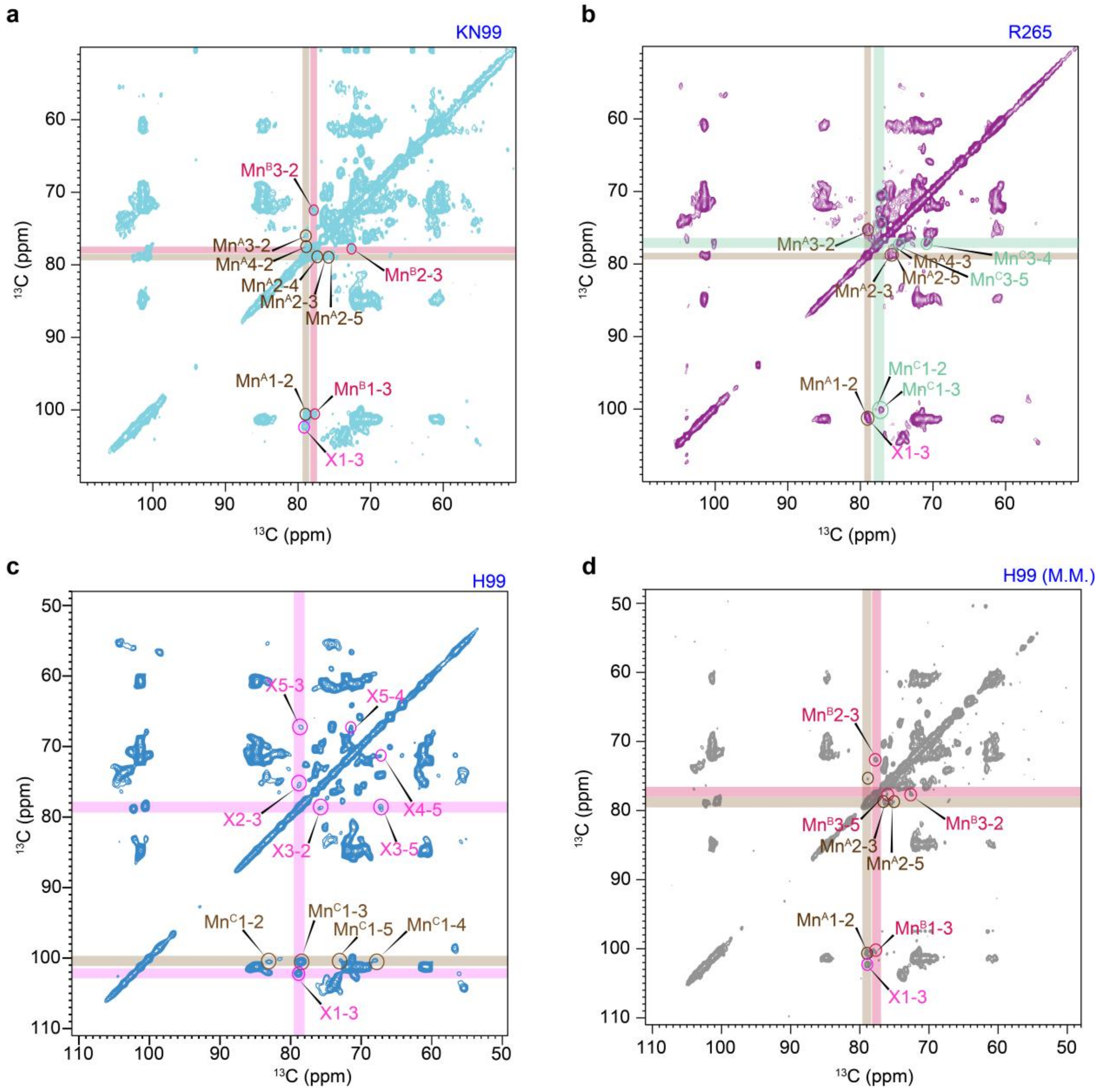


**Supplementary Figure 4. Resonance assignment of capsular molecules in *Cryptococcus*.** CP-based 2D $^{13}C$-$^{13}C$ correlation spectrum measured with 53 ms CORD mixing for different samples of *cryptococcus*, highlighting xylose and different types of mannose signals arising from GXM unit of capsules. **(a)** *C. neoformans* KN99, **(b)** *C. gattii* R265, **(c)** *C. neoformans* H99, and **(d)** *C. neoformans* H99 grown in minimal media.

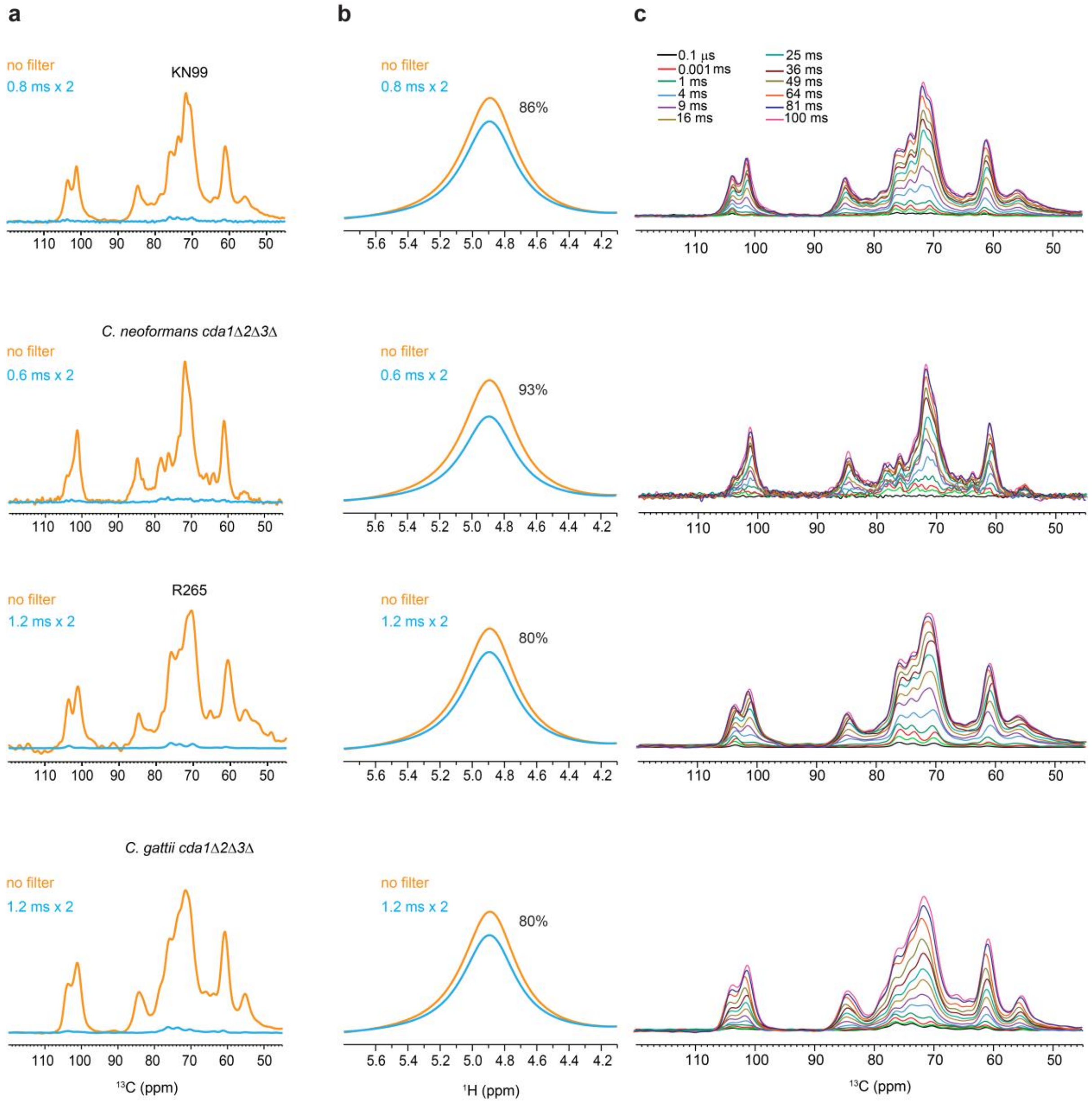


**Supplementary Figure 5. Water-edited experiment setup for inspecting carbohydrate hydration.** **(a)** $^{1}$H-T$_2$ filtered (blue) and control (orange) $^{13}$C spectra are shown for four strains. From the top to bottom: KN99, *C. neoformans cda1Δ2Δ3Δ*, R265, and *C. gattii cda1Δ2Δ3Δ*. No spin diffusion was applied. Approximately 85% of carbohydrate $^{13}$C signals were removed by the T$_2$ filter. **(b)** $^{1}$H-T$_2$ filtered (blue) and control (orange) $^{1}$H NMR spectra, with 80% and 93% of water signal retained for each strain, after the $^{1}$H T$_2$ filter. From the top to bottom: KN99, *C. neoformans cda1Δ2Δ3Δ*, R265, and *C. gattii cda1Δ2Δ3Δ*. **(c)** 1D water-edited $^{13}$C spectra with different $^{1}$H mixing times. From the top to bottom: KN99, *C. neoformans cda1Δ2Δ3Δ*, R265, and *C. gattii cda1Δ2Δ3Δ*. All spectra were measured on 400 MHz spectrometer at 15 kHz MAS at 280 K.

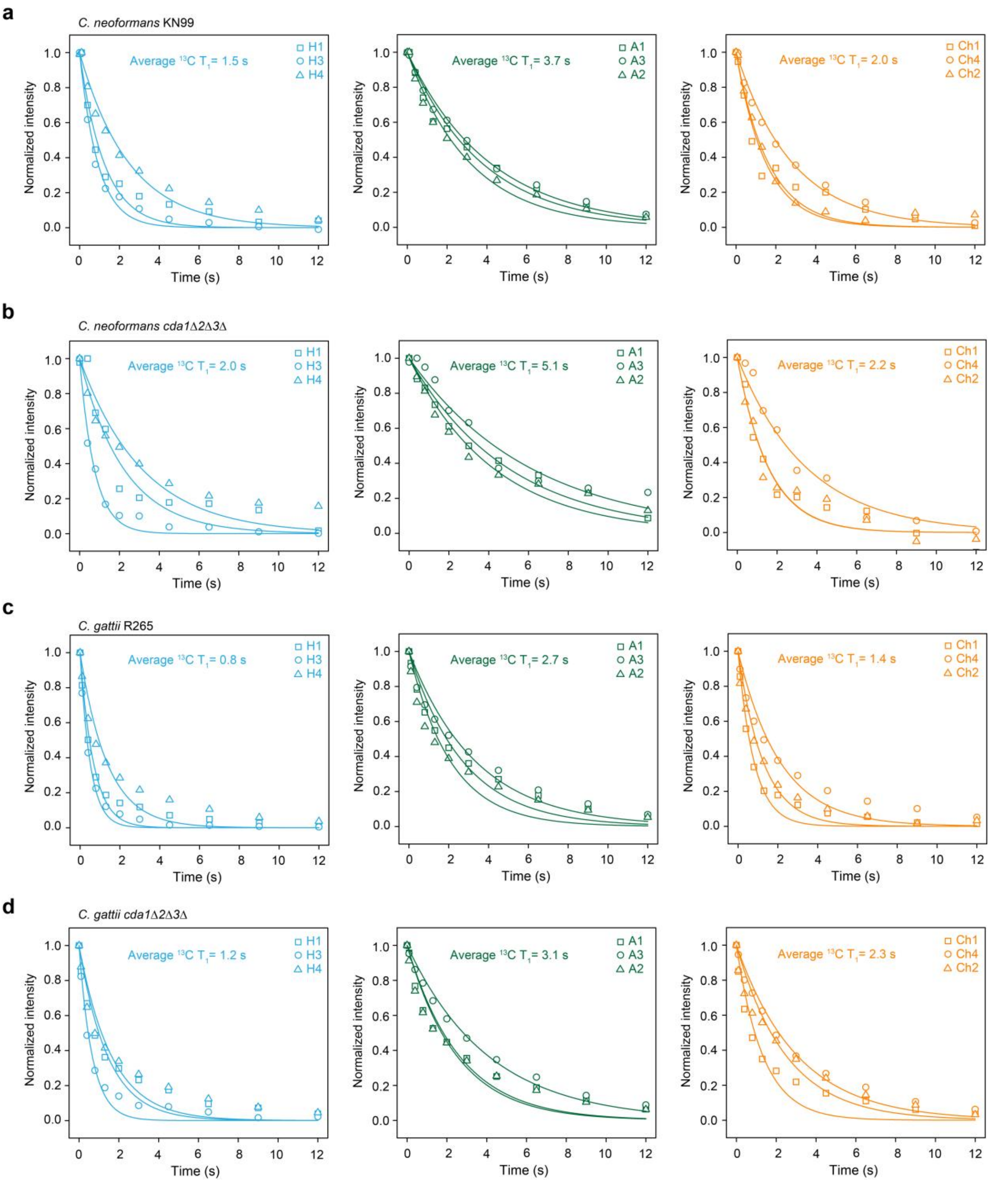


**Supplementary Figure 6. $^{13}$C-T$_1$ relaxation of polysaccharides in *C. neoformans* and *C. gattii*.** $^{13}$C-T$_1$ measured with Torchia CP for **(a)** KN99 **(b)** *C. neoformans cda1Δ2Δ3Δ* **(c)** R265 and **(d)** *C. gattii cda1Δ2Δ3Δ* samples. The data are separately presented for β-1,6-glucan (light blue), α-1,3-glucan (green), and chitin (orange). The acquired data were fitted to a single exponential decay equation. Different symbols and color codes are used to represent different carbons in these polysaccharides.

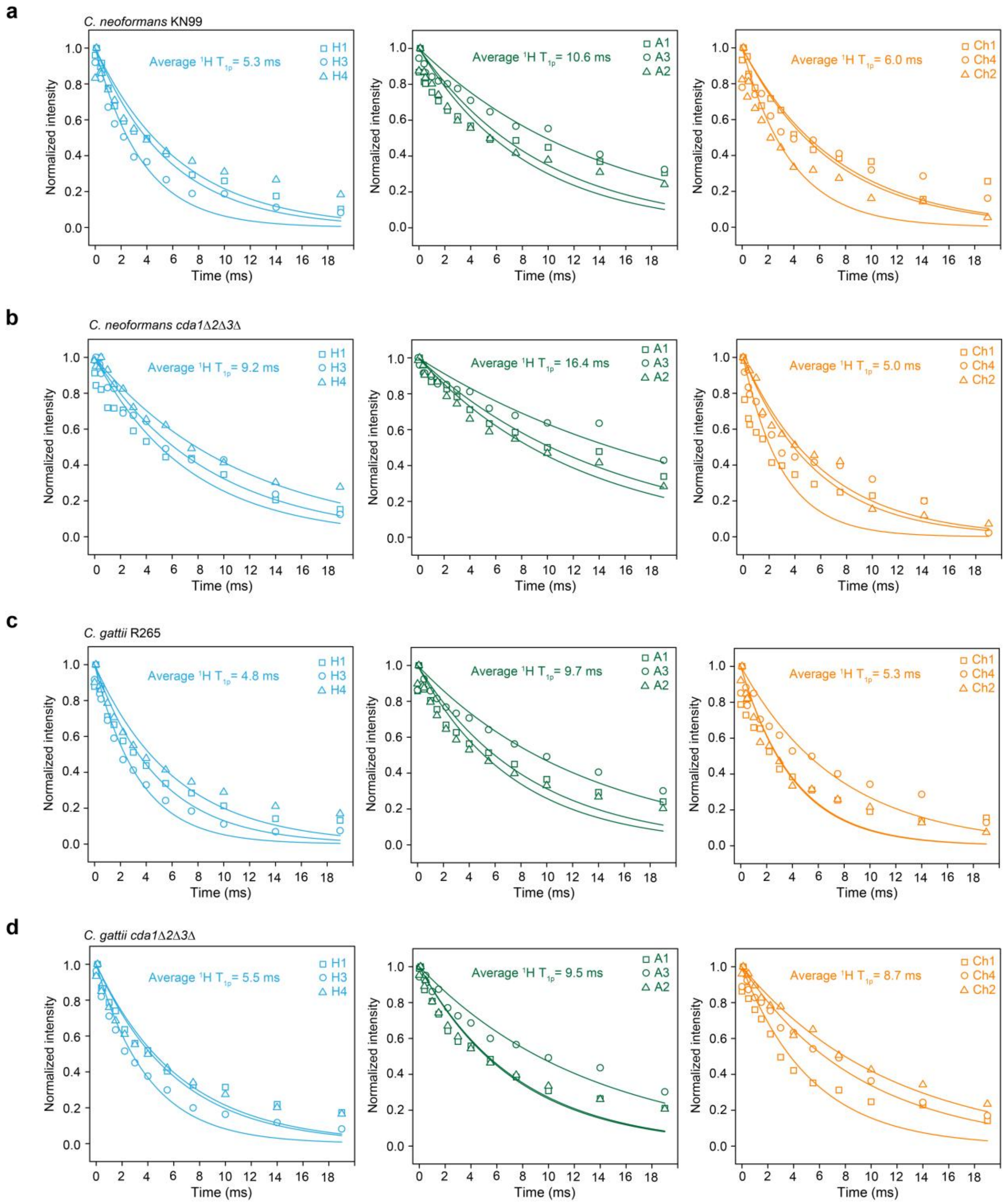


**Supplementary Figure 7. $^{1}$H-T$_{1\rho}$ relaxation of polysaccharides in *C. neoformans* and *C. gattii*.** $^{1}$H-T$_{1\rho}$ measured with Torchia CP for **(a)** KN99 **(b)** *C. neoformans cda1Δ2Δ3Δ* **(c)** R265 and **(d)** *C. gattii cda1Δ2Δ3Δ* samples. The data are separately presented for β-1,6-glucan (light blue), α-1,3-glucan (green), and chitin (orange). The acquired data were fitted to a single exponential decay equation. Different symbols and color codes are used to represent different carbons in these polysaccharides.

**Supplementary Table 1. $^{13}C$ chemical shifts of *C. neoformans* cell wall and capsular polysaccharides in cell walls from $^{13}C$-based experiments**. The referencing scale is TMS scale. All chemical shifts are from room-temperature experiments.

| Carbohydrates | form | C1 | C2 | C3 | C4 | C5 | C6 | Reference |
|---|---|---|---|---|---|---|---|---|
| Rigid molecules | | | | | | | | |
| α-1,3-glucan (A) | a | 101.6 | 72.4 | 85.5 | 69.4 | 71.2 | 61.4 | Chakraborty *et al.* 2021[1] |
| | b | 101.4 | 71.8 | 85.0 | 70.5 | 72.2 | 60.7 | |
| | c | 100.4 | 71.5 | 81.9 | 71.1 | 72.6 | 61.6 | |
| β-1,6-glucan (H) | | 103.8 | 74.2 | 76.8 | 70.8 | 75.8 | 69.8 | Lowman *et al.* 2011[2] |
| Chitin | | 104.3 | 55.3 | 73.7 | 83.7 | 75.7 | 61.1 | Kang *et al.* 2018[3] Fernando *et al.* 2021[4] |
| Chitosan | a | 102.1 | 56.5 | 73.3 | 83.4 | 75.6 | 60.3 | |
| | b | 98.6 | 56.9 | 70.9 | / | / | / | |
| Mannan | Mn[A] | 100.8 | 78.8 | 76.1 | 77.1 | 75.4 | / | Bacon *et al.* 1996[5] Hargett *et al.* 2024[6] Previato *et al.* 2017[7] Ankur *et al.* 2025[8] |
| | Mn[B] | 100.6 | 72.5 | 77.8 | / | 76.0 | / | |
| | Mn[C] | 100.1 | 77.8 | 77.0 | 70.7 | 74.5 | / | |
| Xylose | | 102.2 | 75.8 | 78.8 | 71.3 | 67.1 | n.a. | |
| Mobile molecules | | | | | | | | |
| β-1,6-glucan (H) | | 103.8 | 74.2 | 76.8 | 70.8 | 75.8 | 69.8 | Lowman *et al.* 2011[2] |
| β-1,3-glucan (B) | a | 103.6 | 73.7 | 86.5 | 69.3 | 76.8 | 62.0 | Chakraborty *et al.* 2021[1] Shim *et al.* 2007[9] Fairweather *et al.* 2009[10] Saito *et al.* 1979[11] |
| | b | 103.6 | 73.7 | 85.8 | 69.3 | 76.8 | 62.0 | |
| α-1,3-glucan (A) | a | / | / | / | / | / | / | |
| | b | / | / | / | / | / | / | |
| | c | 100.4 | 71.5 | 81.9 | 71.1 | 72.6 | 61.6 | |
| α-1,2-Mannan (Mn[1,2]) | | 100.7 | 78.7 | 71.3 | 67.2 | 74.0 | 61.7 | Chakraborty *et al.* 2021[1] |
| α-1,6-Mannan (Mn[1,6]) | | 102.9 | 70.8 | / | / | 71.3 | 64.0 | |
| Xylose (X) | | 102.7 | 73.2 | 76.2 | 70.2 | 66.0 | n.a. | Previato *et al.* 2017[7] Ankur *et al.* 2025[8] |

**Supplementary Table 2. The molar composition of rigid polysaccharides.** The numbers are estimated using integrals (volume) of cross peaks in 2D $^{13}C$-$^{13}C$ 53 ms CORD spectra. The average integrals of cross-peaks of each polysaccharide are shown. Error bars are standard errors.

| Strain | α-1,3-glucan | | | β-1,6-glucan | Chitin | Chitosan | | Mannan | Xylose |
|---|---|---|---|---|---|---|---|---|---|
| | a | b | c | | | a | b | | |
| *C. neoformans* KN99 | 36±12 | 26±3 | 9±3 | 15±4 | 3±0.4 | 3±0.4 | 2±0.4 | 2±1 | 4±0.6 |
| *C. neoformans cda1Δ2Δ3Δ* | 42±12 | 28±6 | 6±2 | 7±2 | 7±2 | / | / | 7±4 | 3±1 |
| *C. gattii* R265 | 24±7 | 17±3 | 6±2 | 25±7 | 6±1 | 3±0.3 | 4±0.4 | 9±2 | 6±1 |
| *C. gattii cda1Δ2Δ3Δ* | 24±8 | 16±2 | / | 23±2 | 13±3 | / | / | 10±4 | 14±2 |

(Column group heading printed above the table: Polysaccharide)

The area of the following well-resolved cross peaks 53 ms CORD spectra are used:
α-1,3 (a): the average of C1-C2/3/4 and C3-C2/4.
α-1,3 (b): the average of C1-C2/4, C3-2/4.
α-1,3 (c): the average of C1-C2/4, C3-2/4.
β-1,6: the average of C3-C2/4, C5-C4/6.
Chitin: the average of C1-2/5/6, C4-C2, C5-C2.
Chitosan (a): the average of C1-2/3/5, C3/5-C2.
Chitosan (b): the average of C1-2, C3-C2.
Mannan: the average of C1-C2, C2-C5, C5-4.
Xylose: the average of C1-C3, C2-3.

**Supplementary Table 3. The molar composition of mobile polysaccharides.** The numbers are estimated using integrals (volume) of cross peaks in 2D $^{13}C$-$^{13}C$ refocused DP-J INADEQUATE spectra. The average integrals of cross-peaks of each polysaccharide are shown. Error bars are standard errors of the peak integrals. $Mn^{1,2}$ signals in R265 cells have low intensity.

| | Polysaccharide | | | | | |
|---|---|---|---|---|---|---|
| Strains | β-1,6-glucan | β-1,3-glucan | α-1,3-glucan (c) | Mannan ($Mn^{1,2}$) | Mannan ($Mn^{1,6}$) | Xylose |
| *C. neoformans* KN99 | 79±13 | 8±2 | 2±1 | / | 11±2 | / |
| *C. neoformans cda1Δ2Δ3Δ* | 53±7 | / | 7±1 | 14±2 | 12±2 | 14±2 |
| *C. gattii* R265 | 83±14 | 4±1 | 5±1 | / | 8±3 | / |
| *C. gattii cda1Δ2Δ3Δ* | 74±12 | 8±3 | / | / | 12±3 | 6±1 |

The area of the following well-resolved cross peaks refocused DP-J INADEQUATE spectra are used:
β-1,6: the average of C1, C2, C3, C4, C5, and C6.
β-1,3; the average of C1, C2, C3, C4, C5, and C6.
α-1,3-glucan: the average of C1, C2, and C3.
$Mannan^{1,2}$: the average of C1, C2, C3, C4, C5 and C6.
$Mannan^{1,6}$: the average of C1, C2, C3, C4, C5 and C6.
Xylose: the average of C1, C2, C3, C4, C5 and C6.

**Supplementary Table 4. Water-edited intensities of polysaccharides.** Intensity ratios are obtained by comparing the peak intensities in water-edited and control spectra. The average values for each molecule in each sample are highlighted. Error bars are s.d. propagated from NMR signal-to-noise ratios.

| Polysaccharide | Cross-peak | *C. neoformans* | | *C. gattii* | |
|---|---|---|---|---|---|
| | | KN99 | *cda1Δ2Δ3Δ* | R265 | *cda1Δ2Δ3Δ* |
| α-1,3-glucan (A[a]) | A1-1 | 0.49±0.06 | 0.51±0.03 | 0.33±0.04 | 0.25±0.02 |
| | A1-3 | 0.54±0.08 | 0.55±0.08 | 0.29±0.01 | 0.36±0.06 |
| | A1-2/5 | 0.56±0.02 | 0.47±0.04 | 0.46±0.05 | 0.28±0.03 |
| | A1-A4 | 0.52±0.03 | 0.53±0.06 | 0.28±0.01 | 0.27±0.05 |
| | A3-A1 | 0.32±0.08 | 0.26±0.05 | 0.40±0.02 | 0.34±0.07 |
| | A3-A3 | 0.62±0.05 | 0.36±0.06 | 0.40±0.08 | 0.43±0.05 |
| | A3-2/5 | 0.47±0.06 | 0.59±0.06 | 0.44±0.07 | 0.28±0.04 |
| | A3-4 | 0.36±0.06 | 0.58±0.06 | 0.53±0.02 | 0.21±0.03 |
| | A2/5-1 | 0.49±0.04 | 0.50±0.01 | 0.44±0.04 | 0.20±0.03 |
| | A2/5-3 | 0.54±0.07 | 0.40±0.01 | 0.45±0.08 | 0.22±0.04 |
| | A2/5-2/5 | 0.58±0.01 | 0.53±0.02 | 0.43±0.01 | 0.30±0.01 |
| | A2/5-4 | 0.59±0.09 | 0.45±0.01 | 0.41±0.01 | 0.22±0.02 |
| | A4-1 | 0.49±0.02 | 0.45±0.01 | 0.44±0.02 | 0.56±0.09 |
| | A4-3 | 0.54±0.01 | 0.41±0.01 | 0.52±0.09 | 0.59±0.08 |
| | A4-2/5 | 0.58±0.09 | 0.50±0.03 | 0.48±0.09 | 0.57±0.04 |
| | A4-4 | 0.59±0.03 | 0.53±0.01 | 0.37±0.04 | 0.43±0.01 |
| | Average | 0.52 | 0.48 | 0.42 | 0.34 |
| α-1,3-glucan (A[b]) | A1-1 | 0.49±0.02 | 0.51±0.01 | 0.34±0.04 | 0.25±0.03 |
| | A1-3 | 0.54±0.01 | 0.55±0.06 | 0.29±0.01 | 0.36±0.07 |
| | A1-4 | 0.59±0.06 | 0.52±0.01 | 0.45±0.01 | 0.35±0.04 |
| | A3-A1 | 0.32±0.03 | 0.26±0.01 | 0.39±0.01 | 0.34±0.07 |
| | A3-3 | 0.62±0.06 | 0.36±0.06 | 0.39±0.08 | 0.43±0.05 |
| | A3-4 | 0.49±0.01 | 0.40±0.02 | 0.42±0.06 | 0.25±0.04 |
| | A4-1 | 0.47±0.04 | 0.53±0.06 | 0.49±0.01 | 0.37±0.05 |
| | A4-3 | 0.38±0.06 | 0.42±0.07 | 0.44±0.02 | 0.45±0.05 |
| | A4-4 | 0.54±0.01 | 0.57±0.01 | 0.44±0.08 | 0.35±0.01 |
| | Average | 0.49 | 0.46 | 0.41 | 0.35 |
| α-1,3-glucan (A[c]) | A1-1 | 0.60±0.03 | 0.39±0.01 | 0.34±0.04 | / |
| | A1-3 | 0.78±0.08 | 0.28±0.07 | 0.71±0.02 | |
| | A1-4 | 0.46±0.09 | 0.53±0.01 | 0.49±0.08 | |
| | A3-A1 | 0.23±0.03 | 0.29±0.01 | 0.87±0.04 | |
| | A3-3 | 0.53±0.01 | 0.51±0.03 | 0.53±0.02 | |
| | A3-4 | 0.96±0.04 | 0.66±0.02 | 0.53±0.01 | |
| | A4-1 | 0.22±0.01 | 0.66±0.01 | 0.79±0.09 | |
| | A4-3 | 0.39±0.02 | 0.70±0.01 | 0.60±0.01 | |
| | A4-4 | 0.55±0.02 | 0.54±0.02 | 0.46±0.08 | |
| | Average | 0.52 | 0.51 | 0.59 | |
| β-1,6-glucan (H) | H3-3 | 0.56±0.02 | 0.55±0.04 | 0.48±0.01 | 0.30±0.01 |
| | H3-5 | 0.65±0.02 | 0.60±0.03 | 0.44±0.01 | 0.20±0.02 |
| | H3-2 | 0.58±0.06 | 0.60±0.02 | 0.54±0.03 | 0.26±0.04 |
| | H5-3 | 0.70±0.01 | 0.58±0.01 | 0.23±0.09 | 0.35±0.01 |
| | H5-2 | 0.61±0.03 | 0.63±0.01 | 0.91±0.01 | 0.34±0.02 |
| | H5-5 | 0.31±0.03 | 0.48±0.02 | 0.45±0.04 | 0.17±0.03 |
| | H2-3 | 0.48±0.09 | 0.64±0.04 | 0.46±0.01 | 0.46±0.08 |

| | | | | | |
|---|---|---|---|---|---|
| | H2-5 | 0.76±0.05 | 0.77±0.02 | 0.38±0.05 | 0.40±0.05 |
| | H2-2 | 0.49±0.01 | 0.54±0.01 | 0.44±0.01 | 0.35±0.01 |
| | Average | 0.57 | 0.60 | 0.48 | 0.31 |
| Chitin (Ch) | Ch1-1 | 0.22±0.04 | 0.54±0.01 | 0.36±0.01 | 0.21±0.02 |
| | Ch1-3 | 0.24±0.06 | 0.55±0.02 | 0.67±0.03 | 0.13±0.02 |
| | Ch1-2 | 0.60±0.04 | 0.72±0.02 | 0.88±0.02 | 0.31±0.03 |
| | Ch3-1 | 0.58±0.02 | 0.73±0.03 | 0.76±0.02 | 0.39±0.01 |
| | Ch3-3 | 0.48±0.02 | 0.62±0.02 | 0.47±0.02 | 0.40±0.02 |
| | Ch3-2 | 0.99±0.04 | 0.72±0.08 | 0.43±0.02 | 0.33±0.05 |
| | Ch2-1 | 0.93±0.01 | 0.14±0.02 | 0.94±0.04 | 0.47±0.01 |
| | Ch2-3 | 0.99±0.07 | 0.34±0.03 | 0.49±0.02 | 0.47±0.06 |
| | Ch2-2 | 0.59±0.02 | 0.54±0.01 | 0.34±0.07 | 0.35±0.01 |
| | Average | 0.62 | 0.54 | 0.59 | 0.34 |
| Chitosan (Cs[a]) | Cs1-1 | 0.51±0.05 | / | 0.41±0.05 | / |
| | Cs1-3 | 0.33±0.08 | | 0.62±0.03 | |
| | Cs1-2 | 0.41±0.06 | | 0.68±0.09 | |
| | Cs3-1 | 0.09±0.04 | | 0.97±0.03 | |
| | Cs3-3 | 0.18±0.02 | | 0.47±0.04 | |
| | Cs3-2 | 0.50±0.05 | | 0.53±0.01 | |
| | Cs2-1 | 0.78±0.01 | | 0.63±0.01 | |
| | Cs2-3 | 0.72±0.01 | | 0.44±0.09 | |
| | Cs2-2 | 0.62±0.01 | | 0.42±0.04 | |
| | | 0.48 | | 0.58 | |
| Chitosan (Cs[b]) | Cs1-1 | 0.72±0.05 | / | 0.64±0.05 | / |
| | Cs1-3 | 0.92±0.08 | | 0.85±0.03 | |
| | Cs1-2 | 0.82±0.04 | | 0.72±0.03 | |
| | Cs3-1 | 0.40±0.06 | | 0.80±0.09 | |
| | Cs3-3 | 0.58±0.05 | | 0.45±0.04 | |
| | Cs3-2 | 0.47±0.02 | | 0.54±0.01 | |
| | Cs2-1 | 0.76±0.01 | | 0.97±0.01 | |
| | Cs2-3 | 0.98±0.01 | | 0.57±0.04 | |
| | Cs2-2 | 0.58±0.01 | | 0.48±0.08 | |
| | | 0.69 | | 0.67 | |

**Supplementary Table 5. $^{1}H\text{-}T_{1\rho}$ and $^{13}C\text{-}T_1$ relaxation times of polysaccharides in cell walls.** Data is shown for the *C. neoformans* samples. The average values for each molecule in each sample are highlighted in bold. The data were measured using 1D $^{13}C$ relaxation experiments. The data are fit using single exponential equations: $I(t) = e^{-t/T_1}$. Error bars are standard deviations of the fit parameters.

| Polysaccharide | Chemical shift | KN99 | | *cda1Δ2Δ3Δ* | | R265 | | *cda1Δ2Δ3Δ* | |
|---|---|---|---|---|---|---|---|---|---|
| | | $^{1}H\text{-}T_{1\rho}$ (ms) | $^{13}C\text{-}T_1$ (s) | $^{1}H\text{-}T_{1\rho}$ (ms) | $^{13}C\text{-}T_1$ (s) | $^{1}H\text{-}T_{1\rho}$ (ms) | $^{13}C\text{-}T_1$ (s) | $^{1}H\text{-}T_{1\rho}$ (ms) | $^{13}C\text{-}T_1$ (s) |
| α-1,3-glucan (a) | 101.5 | 9.5±1.4 | 3.7±0.2 | 14.8±0.9 | 5.0±0.3 | 8.6±0.8 | 3.4±0.3 | 7.5±0.6 | 2.6±0.2 |
| | 85.0 | 14.2±1.0 | 4.1±0.1 | 21.7±1.4 | 6.2±0.5 | 7.4±0.7 | 2.7±0.2 | 7.6±0.6 | 2.5±0.2 |
| | 71.9 | 8.3±0.8 | 3.1±0.2 | 12.6±0.7 | 4.3±0.3 | 13.2±1.0 | 2.1±0.2 | 13.3±0.9 | 4.1±0.2 |
| | Average | 10.6 | 3.7 | 16.4 | 5.1 | 9.7 | 2.7 | 9.5 | 3.1 |
| Chitin | 104.6 | 7.4±0.7 | 1.5±0.2 | 3.0±0.5 | 1.6±0.2 | 4.0±0.5 | 1.3±0.1 | 5.4±0.5 | 1.3±0.2 |
| | 83.4 | 7.0±0.9 | 2.8±0.1 | 5.6±0.6 | 3.5±0.3 | 4.1±0.4 | 0.8±0.07 | 11.5±0.5 | 2.5±0.2 |
| | 55.9 | 3.8±0.4 | 1.6±0.1 | 6.2±0.4 | 1.6±0.2 | 7.6±0.8 | 2.1±0.2 | 9.2±0.6 | 3.1±0.1 |
| | Average | 6.0 | 2.0 | 5.0 | 2.2 | 5.3 | 1.4 | 8.7 | 2.3 |
| β-1,6-glucan | 103.9 | 5.7±0.4 | 1.2±0.1 | 7.3±0.7 | 2.2±0.3 | 3.4±0.2 | 0.7±0.06 | 6.1±0.5 | 1.6±0.2 |
| | 76.4 | 3.6±0.3 | 0.9±0.07 | 8.8±0.5 | 0.7±0.05 | 4.9±0.4 | 0.5±0.03 | 6.5±0.5 | 1.4±0.1 |
| | 70.6 | 6.5±0.7 | 2.5±0.2 | 11.4±0.5 | 3.1±0.4 | 6.2±0.5 | 1.4±0.2 | 4.0±0.3 | 0.6±0.06 |
| | Average | 5.3 | 1.5 | 9.2 | 2.0 | 4.8 | 0.8 | 5.5 | 1.2 |

**Supplementary Table 6. $^{13}$C Solid-state NMR experimental parameters for fungal cell wall characterization.** T = sample temperature; $B_0$ = magnetic field; $\nu_{MAS}$ = MAS frequency; ns = number of scans; $d_1$ = recycle delay between scans; $t_{1,\,max}$ = maximum $t_1$ evolution time (for indirect dimension); $t_{1,\,inc}$ = increment for $t_1$ (for indirect dimension) evolution time; $\tau_{dw}$ = dwell time during direct FID acquisition; $\tau_{acq}$ = maximum acquisition time during direct FID detection; $\tau_{XY}$ = cross-polarization contact time during CP from channel X to channel Y; $\nu_{1H,\,dec}$ = dipolar decoupling field strength**.** Spin diffusion (SD).

| Experiment | NMR Parameters | | | | | | | | | | | | | Samples |
|---|---|---|---|---|---|---|---|---|---|---|---|---|---|---|
| | T (K) | $B_0$ (T) | $\nu_{MAS}$ (kHz) | ns | d1 (s) | $t_{1,\,max}$ (ms) | $t_{1,\,inc}$ (μs) | $\tau_{dw}$ (μs) | $\tau_{acq}$ (ms) | $\tau_{HC}$ (ms) | $\tau_{SD}$ (ms) | $\tau_{mix}$ (ms) | $\nu_{1H\ dec}$ (kHz) | |
| Identification and quantification of polysaccharides | | | | | | | | | | | | | | KN99, R265, *cda1Δ2Δ3Δ* |
| 1D $^{13}$C CP | 298 | 18.8 | 15 | 1024 | 2 | | | 5 | 18 | 1 | | | 83 | |
| 1D $^{13}$C DP | 298 | 18.8 | 15 | 512 | 2 or 35 | | | 5 | 18 | | | | 83 | |
| 2D $^{13}$C-$^{13}$C with CORD mixing | 298 | 18.8 | 15 | 32 | 2 | 7.5 | 25 | 5 | 14 | 1 | | 53 $\tau_{CORD}$ | 83 | |
| 2D $^{13}$C-$^{13}$C refocused CP J-INADEQUATE | 298 | 18.8 | 15 | 16 | 2 | 7.5 | 22 | 5 | 14 | | | | 83 | |
| 2D $^{13}$C-$^{13}$C refocused DP J-INADEQUATE | 298 | 18.8 | 15 | 16 | 2 | 7.5 | 22 | 5 | 14 | | | | 83 | |
| PAR | 280 | 18.8 | 15 | 32 | 2 | 7.5 | 20 | 5 | 20.5 | 1 | | 15 | 83 | R265, *cda1Δ2Δ3Δ* |
| Estimation of site-specific hydration of polysaccharides | | | | | | | | | | | | | | KN99, R265, *cda1Δ2Δ3Δ* |
| 2D $^{13}$C-$^{13}$C water-edited | 280 | 9.4 | 15 | 64 | 2 | 5.5 | 50 | 8 | 16 | 1 | 0, 4 | 50 $\tau_{PDSD}$ | 71 | |
| Dynamics of polysaccharides | | | | | | | | | | | | | | |
| 1D $^{13}$C-$T_1$ | 298 | 9.4 | 15 | 512 | 2 | | | 8 | 16 | 1 | | | 71 | |
| 1D $^{1}$H-$T_{1\rho}$ | 298 | 9.4 | 15 | 512 | 2 | | | 8 | 16 | 1 | | | 71 | |
